# Small-molecule targeting of GPCR-independent non-canonical G protein signaling inhibits cancer progression

**DOI:** 10.1101/2023.02.18.529092

**Authors:** Jingyi Zhao, Vincent DiGiacomo, Mariola Ferreras-Gutierrez, Shiva Dastjerdi, Alain Ibáñez de Opakua, Jong-Chan Park, Alex Luebbers, Qingyan Chen, Aaron Beeler, Francisco J Blanco, Mikel Garcia-Marcos

## Abstract

Activation of heterotrimeric G-proteins (Gαβγ) by G-protein-coupled receptors (GPCRs) is a quintessential mechanism of cell signaling widely targeted by clinically-approved drugs. However, it has become evident that heterotrimeric G-proteins can also be activated via GPCR-independent mechanisms that remain untapped as pharmacological targets. GIV/Girdin has emerged as a prototypical non-GPCR activator of G proteins that promotes cancer metastasis. Here, we introduce IGGi-11, a first-in-class smallmolecule inhibitor of non-canonical activation of heterotrimeric G-protein signaling. IGGi-11 binding to G-protein α-subunits (Gαi) specifically disrupted their engagement with GIV/Girdin, thereby blocking non-canonical G-protein signaling in tumor cells, and inhibiting pro-invasive traits of metastatic cancer cells *in vitro* and in mice. In contrast, IGGi-11 did not interfere with canonical G-protein signaling mechanisms triggered by GPCRs. By revealing that small molecules can selectively disable non-canonical mechanisms of G-protein activation dysregulated in disease, these findings warrant the exploration of therapeutic modalities in G-protein signaling that go beyond targeting GPCRs.

## Introduction

G protein-coupled receptors (GPCRs) mediate a large fraction of all transmembrane signaling in the human body, including responses triggered by every major neurotransmitter and by two-thirds of hormones (1). They are also the largest family of druggable proteins in the human genome, representing the target for over one-third of clinically approved drugs (2). To relay signals, GPCRs activate heterotrimeric G-proteins (Gαβγ) in the cytoplasm by promoting the exchange of GDP for GTP on Gα subunits, which results in a concomitant dissociation of Gβγ dimers (3). In turn, Gα-GTP and “free” Gβγ act on downstream effectors to propagate signaling. Signaling is turned off by the intrinsic GTPase activity of Gα, which leads to the re-association of Gα with Gβγ. There is also a growing number of cytoplasmic proteins that modulate nucleotide handling by G-proteins, thereby exerting profound effects on the duration and amplitude of signaling (4–11).

In stark contrast to GPCRs, there are no clinically approved drugs for heterotrimeric G-proteins, despite their well-documented potential as pharmacological targets (12). Small molecule inhibitors of Gβγ have been validated in some preclinical models (12, 13), but no drug-like small molecule that targets Gα subunits has been validated. There are, however, some natural cyclic depsipeptides that block α-subunits of the Gq/11 family with high specificity and potency (14). Unfortunately, because they inhibit G-protein activation *en toto*, these compounds could cause undesired side effects due to indiscriminate blockade of ubiquitous, physiologically-relevant functions of their target G-proteins.

Perhaps a more nuanced targeting approach that exploits disease-specific mechanisms of G-protein regulation could pave the way for new pharmacology. This idea is thwarted by the realization that the mechanisms of G-protein regulation beyond ubiquitous GPCR-mediated activation remain poorly understood in the absence of adequate tools to interrogate them. GIV (also known as Girdin) is a cytoplasmic protein that binds to Gαi subunits to promote G-protein signaling in a GPCR-independent manner (8, 15–17) and its expression in human primary solid tumors correlates with progression towards more invasive, metastatic stages in various types of cancer (18–20). Tumor cells depleted of GIV also fail to migrate *in vitro* or metastasize in mice (21). Here, we report the identification of a small molecule that binds to Gαi to selectively prevent GIV binding without disturbing other mechanisms by which the G-protein is regulated, including canonical GPCR-mediated signaling. We leverage this compound to establish that GIV-mediated activation of G-protein signaling favors cancer progression by operating downstream of receptor tyrosine kinases (RTKs) instead of downstream of GPCRs.

## Results

### High-throughput screen for inhibitors of the GIV-Gαi interaction

Previous work indicates that expression of GIV at high levels in cancer cells might facilitate its association with Gαi, which in turn favors tumor cell migration and other pro-metastatic traits (8, 15–17, 22–25) (**Fig. 1A**). Moreover, characterization of the molecular basis for the GIV-Gαi interaction (**Fig. 1A**) revealed that this protein-protein interaction might be suitable for specific pharmacological disruption (26–28). These previous findings motivated us to pursue a small molecule screen for inhibitors of the GIV-Gαi interaction. Using a fluorescence polarization (FP) assay that directly monitors GIV binding to Gαi3 (27), we obtained 580 hits from screening a collection of 200,000 compounds (**Fig. 1B, C**). Of these, 155 tested positive for inhibition in both the primary FP assay and an orthogonal secondary assay (AlphaScreen^®^, AS) (27) (**Fig. 1C, D**). After triage, 68 compounds were discarded based on unfavorable chemical properties, and only 69 of the remaining 87 compounds could be repurchased as fresh powder stocks (**Fig. 1D, Table S1**). We named this set of compounds “IGGi”, for “Inhibitors of the GIV-Gαi interaction”. We next evaluated the performance of these 69 IGGi compounds in cell-based assays. In cancer cell lines that express high levels of GIV (e.g., the triple-negative metastatic breast cancer cell line, MDA-MB-231), loss of GIV or disruption of its ability to bind Gαi through mutagenesis impairs cell migration, but does not affect cell viability under standard *in vitro* culture conditions on plastic dishes (17, 21). We found that approximately one-third of the IGGi compounds impaired MDA-MB-231 cell migration without affecting viability (**Fig. 1E**), lending confidence on the ability of our biochemical screen to identify compounds with the desired biological activity. To further prioritize the 69 IGGi compounds, we excluded not only those with the undesired property of reducing MDA-MB-231 viability, but also those that reduced the viability of MCF-7 cells (a non-metastatic breast cancer cell line that expresses low levels of GIV) or of MCF-10A (a non-transformed epithelial breast cell line) to eliminate molecules with non-specific cytotoxicity (**Fig. 1F**). The remaining 44 compounds were tested in a tertiary GIV-Gαi binding assay based on GST-fusion pull-downs (PD) (**Fig. 2A**). Only one compound, IGGi-11, was found to inhibit Gαi3 binding to GIV in this assay. Despite the weak activity of this compound in MDA-MB-231 cell migration assays (**Fig. 1E**), experiments presented below indicated high specificity and suitability for cell-based systems upon analog development.

**Figure 1.**
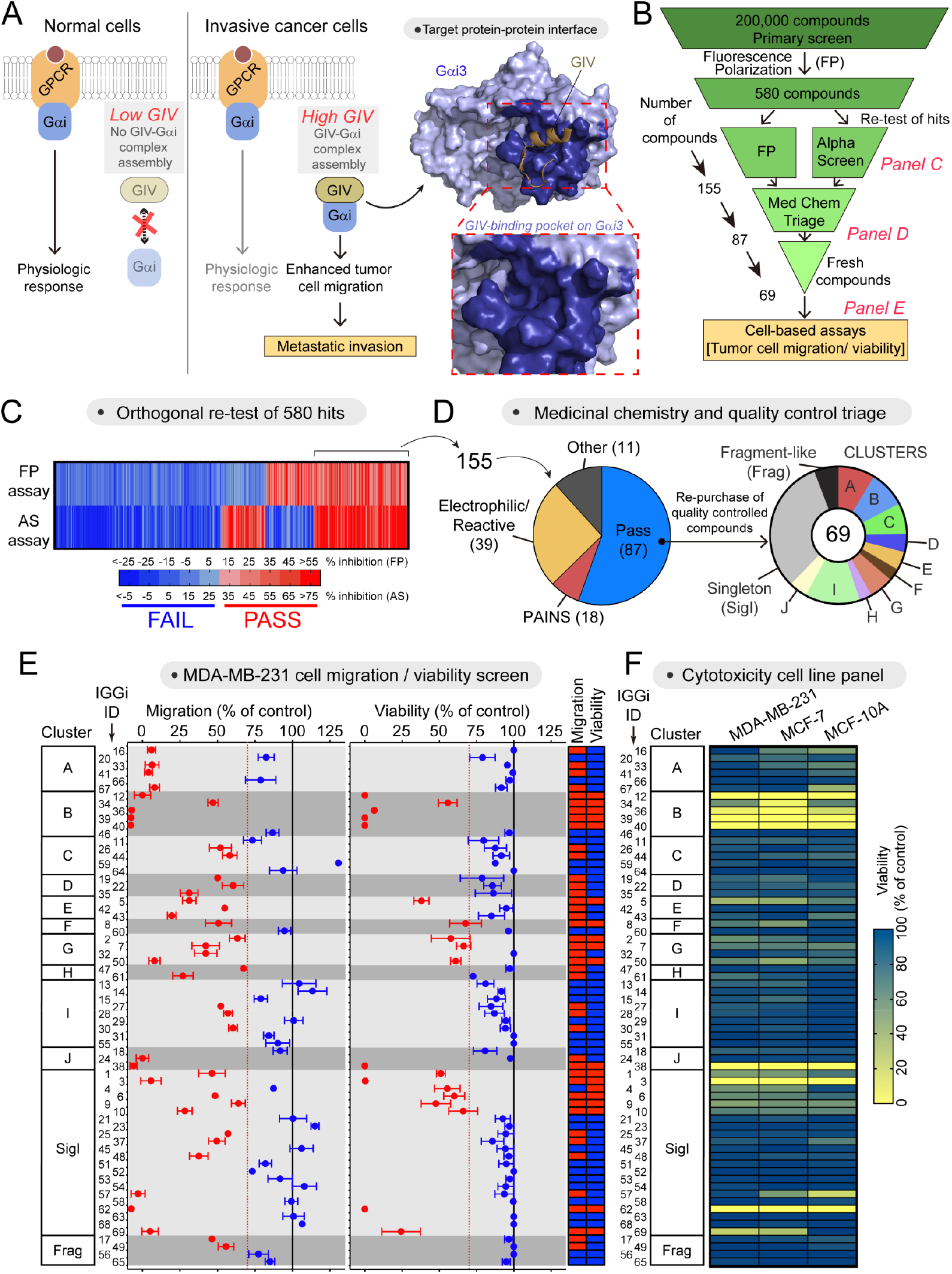
Small molecule screening to identify inhibitors of the GIV-Gαi interaction. **(A)** Diagram depicting the rationale for targeting the GIV-Gαi interaction with small molecules. **(B)** Scheme of the full screening campaign. **(C)** Confirmation of hit compounds that inhibit the GIV-Gαi interaction in two orthogonal biochemical assays, fluorescence polarization (FP) and AlphaScreen (AS). **(D)** Triage of compounds based on unfavorable chemical properties and availability of molecules with quality control. **(E)** Test of 69 IGGi compounds (100 μM) on MDA-MB-231 cell migration and viability. Red, <30% reduction; blue, >30% reduction. Mean ± SEM (*N* = 4). **(F)** Comparison of the effect of IGGi compounds (100 μM) on the viability of three breast cell lines, MDA-MB-231, MCF-7, and MCF-10A (mean of *N* = 3).

### IGGi-11 binds to the GIV interacting region of Gαi

We reasoned that inhibitors of the GIV-Gαi interaction should bind to the G-protein because our primary screening assay used a small peptide fragment of GIV unlikely to harbor enough structural features to accommodate a small molecule. Using nuclear magnetic resonance (NMR) spectroscopy, we found that IGGi-11 caused dose-dependent chemical shift perturbations (CSP) in the amide bond signals of discrete amino acids of isotopically labeled (^2^H-^13^C-^15^N) Gαi3 (**Fig. 2B**, **Fig. S1**), indicating compound binding. In contrast, another IGGi compound, IGGi-41, that was a potent inhibitor of MDA-MB-231 cell migration (**Fig. 1E**) but did not disrupt GIV-Gαi binding (**Fig. 2A**), did not cause NMR signal perturbations (**Fig. S2**). These results suggested that IGGi-11 binds specifically to Gαi3. When IGGi-11-induced NMR perturbations were overlaid on a structural model of IGGi-11-bound Gαi3 and compared to a structural model of the GIV-Gαi3 complex, several of the amino acids with the largest perturbations (S252, W258, F259, F215, E216, G217, and K35) clustered around the predicted docking site for IGGi-11 and overlapped with the binding area for GIV (**Fig. 2C**). To directly test if IGGi-11 binds on this predicted site located in the groove between the α3 helix and the conformationally dynamic Switch II (SwII) region, we carried out isothermal titration calorimetry (ITC) experiments with wild-type Gαi3 (WT) or mutants. We found that three different mutations in the predicted binding site for IGGi-11 on Gαi3 (F215A, N256E, and W258A) lead to large decreases in compound binding affinity (>10-30 fold), whereas another mutation in an amino acid adjacent to the predicted binding site (G42R) did not (**Fig. 2D**). All mutant proteins fold properly and remain functional based on multiple assays (26). The estimated equilibrium dissociation constant (K_D_) for the Gαi3/IGGi-11 interaction based on ITC was ~4 μM (**Fig. 2D**), which was in good agreement with estimates based on curve fits of CSPs observed in NMR experiments (0.9-4.6 μM, **Fig. S1**). IGGi-11 also blocked GIV binding to Gαi3 in FP assays with an inhibition constant (K_i_) of ~14 μM, and similar results were obtained for the other two Gα proteins of Gi family: Gαi1 and Gαi2 (**Fig. S3A**). Consistently, IGGi-11 also inhibited the ability of GIV to promote the steady-state GTPase activity of Gαi3, which reports increased nucleotide exchange *in vitro* (25) (**Fig. S3B**). Together, these results indicate that IGGi-11 binds to the GIV interacting site of Gαi proteins with low micromolar affinity, thereby precluding the formation of the GIV-Gαi complex *in vitro*.

**Figure 2.**
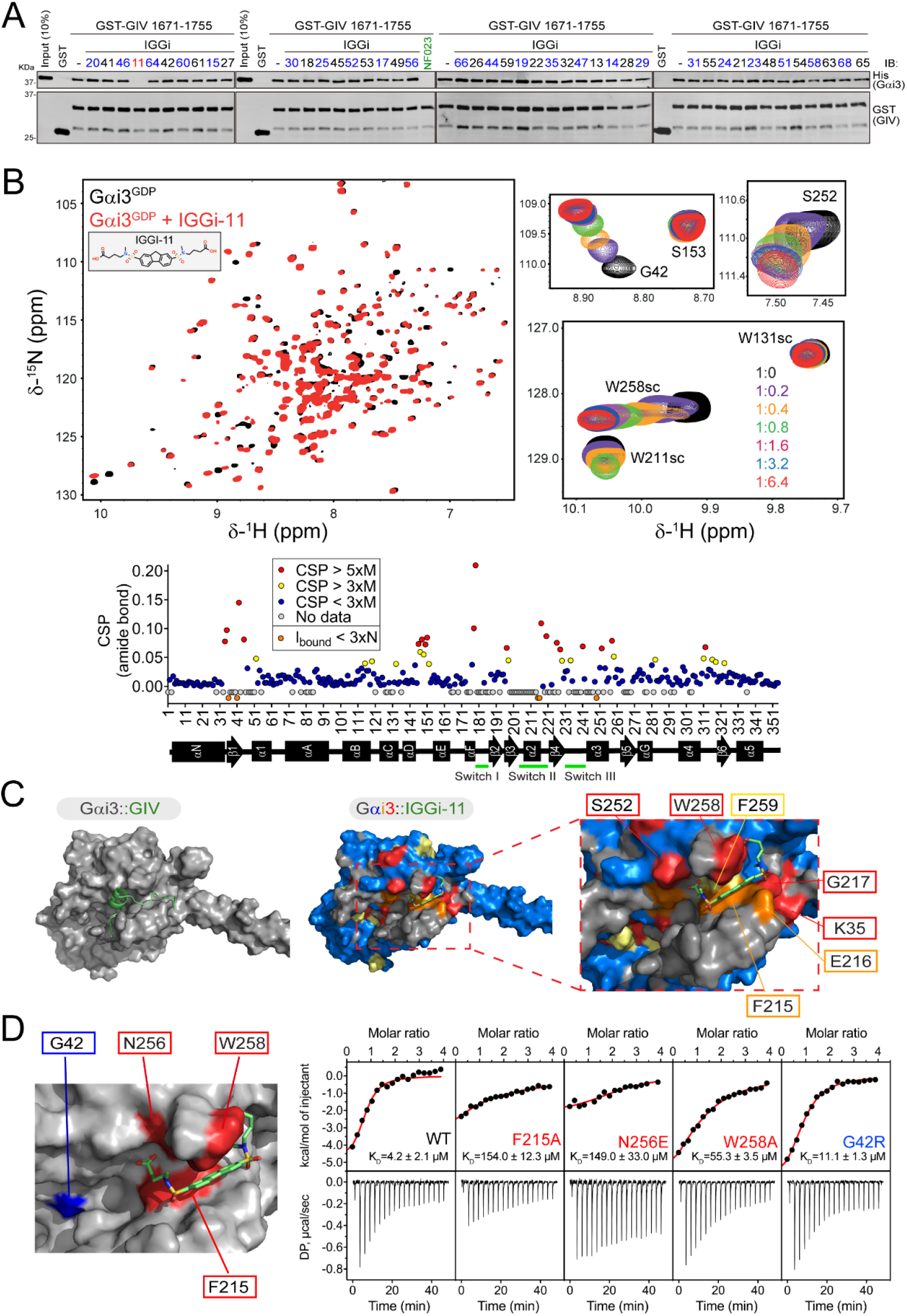
IGGi-11 binding to the GIV interacting region of Gαi. **(A)** IGGi-11 disrupts GIV-Gαi binding in pulldown assays. His-Gαi3 was incubated with glutathione agarose-bound GST-GIV (aa 1671-1755) in the presence of the indicated compounds or the positive control NF023 at a concentration of 100 μM. After incubation and washes, bead-bound proteins were separated by SDS-PAGE and immunoblotted (IB) as indicated. Representative of 3 independent experiments. **(B)** Overlay of ^1^H-^15^N TROSY spectra of ^2^H,^13^C,^15^N-Gαi3-GDP in the absence or presence of IGGi-11. Selected regions from the overlaid spectra depicting representative perturbations in Gαi3 signals induced by increasing amounts of IGGI-11 are shown at the right. The dot plot (bottom) corresponds to the quantification of IGGi-11 induced chemical shift perturbations (CSPs). Red, CSP > 5 times the median (M); yellow, CSP >3xM; blue, CSP <3xM; grey, no data. Reductions in signal intensity (I_bound_) below 3 times the noise (N) are indicated in orange. **(C)** Comparison of models of IGGi-11 docked onto Gαi3 (*middle and right*, color coded according to NMR perturbations quantified in panel A) and GIV-bound Gαi3 (*left*). **(D)** Quantification of IGGi-11 binding affinity (K_D_) for Gαi3 wild-type (WT) or the indicated mutants using isothermal titration calorimetry (ITC). Data are representative of at least two independent experiments.

### IGGi-11 does not affect GIV-independent aspects of G-protein regulation and function

A concern with targeting Gαi is the potential on-target but nonetheless undesired effects that may result due to the many functions of G-proteins. The activity of Gα subunits depends on the ability to handle nucleotides (GDP/GTP exchange, GTP hydrolysis), on proteins that regulate their activity (Gβγ, GPCRs, Guanine nucleotide Dissociation Inhibitors (GDIs), and GTPase Accelerating Proteins (GAPs)), or on how they regulate other proteins that propagate signaling (effectors) (**Fig. 3A**). With this in mind, we set out to thoroughly address the potential effect of IGGi-11 on G-protein functions other than those mediated via GIV binding by using isolated cell membranes or purified proteins. First, we tested the effect of IGGi-11 on the association of Gβγ with Gα using a bioluminescence resonance energy transfer (BRET) assay (29, 30). We found that concentrations of IGGi-11 up to 100 μM did not cause the dissociation of Gβγ from Gαi3 (**Fig. 3B**), whereas incubation with a positive control peptide (R12 GL, 25 μM) or a non-hydrolyzable GTP analog (GTPγS, 300 μM) did. Similar observations were made with three other Gα subunits that belong to the same family as Gαi (i.e., Gαo), or to different ones (i.e., Gαq and Gα13) (**Fig. S4A**), indicating that IGGi-11 does not disrupt Gαβγ heterotrimers. Using the same assay, we assessed the effect of IGGi-11 on GPCR-mediated activation of G-proteins, which results in the dissociation of Gβγ from Gα. We found that concentrations of IGGi-11 up to 100 μM did not interfere with the ability of agonist-stimulated GPCRs to activate Gi3, Go, Gq, or G13 heterotrimers (**Fig. 3C**, **Fig, S4B**). Rapid kinetic assays further confirmed that IGGi-11 did not alter the rate of Gβγ dissociation upon GPCR activation (**Fig. 3D**) or the rate of Gβγ-Gαi3 reassociation upon GPCR signal termination (**Fig. S4C**). As an alternative to assess GPCR-mediated activation of G-proteins, we used another BRET-based biosensor (31) that directly monitors the formation of GTP-bound Gαi3 (**Fig. 3E**). We found that neither amplitude nor kinetics of Gαi3-GTP formation upon GPCR stimulation were affected by IGGi-11 (**Fig. 3E, F**). We also found that IGGi-11 did not interfere with the spontaneous exchange of GDP for GTP on Gαi3 using three independent assays: BRET-based GTPγS binding to Gαi in isolated membranes (**Fig. S5A**), binding of fluorescently labeled GTPγS to purified Gαi (**Fig. S5B**), or steady-state GTPase activity of purified Gαi with radiolabeled GTP (**Fig. S5C**). We also found that IGGi-11 did not affect the hydrolysis of GTP to GDP by purified Gαi (**Fig. S5D**).

**Figure 3.**
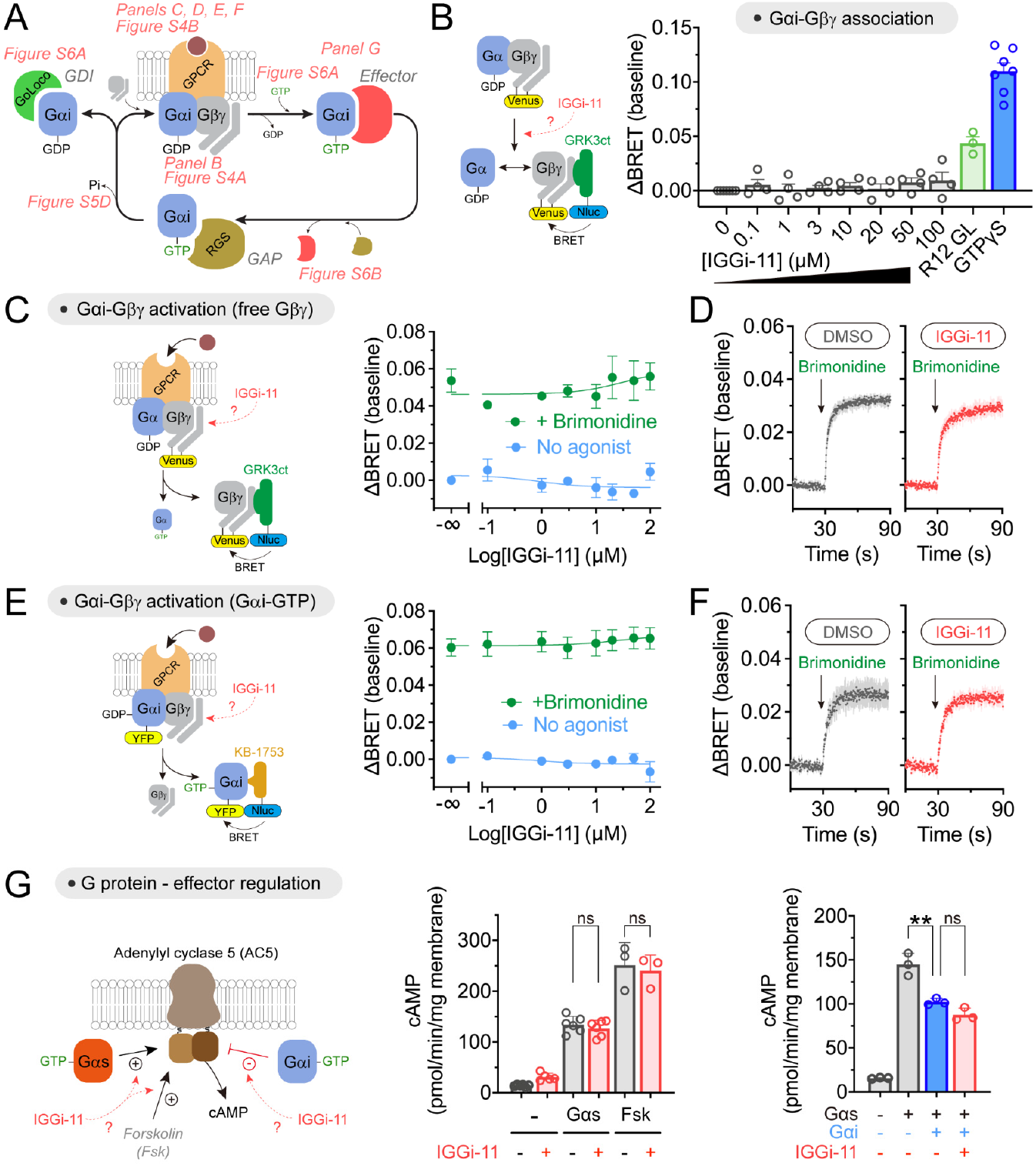
Lack of effect of IGGi-11 on G-protein coupling to GPCRs and effectors. **(A)** Diagram of key steps and protein interactions involved in Gαi-subunit functions. **(B)** IGGi-11 does not dissociate Gβγ from Gαi3 in membranes isolated from HEK293T cells expressing a BRET-based biosensor for free Gβγ, whereas two positive controls do (a GoLoco peptide derived from RGS12, R12 GL, 25 μM; and GTPγS 300 μM). **(C-F)** IGGi-11 does not affect GPCR-mediated activation of Gi3 as determined by the dissociation of Gαi3-Gβγ heterotrimers (C, D) or the formation of Gαi3-GTP (E, F) using BRET-based biosensors. In C and E, membranes isolated from HEK293T cells expressing the α2_A_ adrenergic receptor were treated with the indicated concentrations of IGGi-11 with (green) or without (blue) stimulation with a receptor agonist (brimonidine, 1 μM) for 2 minutes before BRET measurements. In D and F, BRET was continuously measured in real time in the presence of 100 μM IGGi-11 or vehicle (1% DMSO, v:v). **(G)** IGGi-11 does not interfere with G-protein-mediated regulation of adenylyl cyclase. Membranes isolated from HEK293T cells expressing adenylyl cyclase 5 were treated with IGGi-11 (100 μM), purified Gαs (0.5 μM), purified myristoylated Gαi1 (Gαi, 1 μM), and forskolin (Fsk, 10 μM) in the combinations indicated in the graphs. Mean ± SEM (*N* ≥ 3). *\*\*P* ≥ 0.01, ANOVA.

Next, we evaluated the potential impact of IGGi-11 on the ability of active, GTP-bound Gαi proteins to engage and modulate effectors. First, we observed that IGGi-11 did not cause NMR signal perturbations in the α3/SwII region of GTPγS-loaded Gαi3 (**Fig. S6A**), which contrasts with the observations obtained for GDP-loaded Gαi3 (**Fig. 2B**, **Fig. S1**) and suggests lack of compound binding to active G-proteins. Consistent with this, we also found that IGGi-11 did not inhibit the interaction between purified Gαi3 and KB-1753, an effector-like peptide that binds to the α3/SwII region of Gαi-GTP (32) (**Fig. S6B**). We then tested whether IGGi-11 affected the regulation of a *bona fide* effector of Gαi, i.e., adenylyl cyclase (**Fig. 3G**). In membranes from cells expressing adenylyl cyclase 5, IGGi-11 did not affect either activation mediated by purified Gαs or inhibition mediated by purified Gαi (**Fig. 3G**). The compound did not affect adenylyl cyclase activity either under basal conditions or upon direct, G-protein-independent activation with forskolin (**Fig. 3G**).

Finally, we assessed whether IGGi-11 would preclude the binding of Gαi to other G-protein regulators like Guanine nucleotide Dissociation Inhibitors (GDIs) that contain a GoLoco motif (4, 5), or GTPase-Accelerating Proteins (GAPs) of the Regulators of G protein Signaling (RGS) family (6, 7). We found that IGGi-11 did not inhibit the interaction of Gαi3 with the GoLoco motif responsible for the GDI activity of RGS12 (R12 GL, **Fig. S6B**) or with the GAP RGS4 (**Fig. S6C**).

Taken together, our results indicate that IGGi-11 specifically inhibits GIV binding to Gαi without interfering with any other major function of Gαi, including nucleotide binding and hydrolysis, association with Gβγ subunits and other cytoplasmic regulators, activation by GPCRs, or modulation of effectors.

### Validation of an IGGi-11 analog with increased activity in cells

After establishing the specificity of IGGi-11 for the target GIV-Gαi complex *in vitro*, we sought to determine its biological activity in cells. We found that preincubation of MDA-MB-231 cells with IGGi-11 inhibited their ability to migrate only marginally (**Fig. S7A**). We reasoned that this could be due to low membrane permeability because IGGi-11 contains two negatively charged carboxylate groups (**Fig. S7A**). To overcome this, we generated IGGi-11me, an analog in which the carboxylates were esterified with methyl groups. We hypothesized that esterification would increase membrane permeability by eliminating the charges of the carboxylates, and that cytoplasmic esterases would revert the modification to produce IGGi-11, thereby enabling enhanced inhibitory activity in cells (**Fig. S7A**). Indeed, preincubation of MDA-MD-231 cells with IGGi-11me efficiently reduced their ability to migrate (**Fig. S7A**). As desired, IGGi-11me (or IGGi-11) did not affect the viability of MDA-MB-231 or MCF-10A cells (**Fig. S7B**). We confirmed that IGGi-11me had higher permeability than IGGi-11 by using a parallel artificial membrane permeability assay (**Fig. S7C**). We also confirmed that IGGi-11me was converted to IGGi-11 by esterases present in the cytosol of MDA-MB-231 cells (**Fig. S7D**), which is a critical step because IGGi-11me inhibits GIV-Gαi3 binding with lower potency than IGGi-11 (**Fig. S7E**). These results indicate that IGGi-11me serves as pro-drug that allows the action of the active GIV-Gαi inhibitor compound, IGGi-11, in cells.

### IGGi-11me inhibits GIV-dependent cancer cell signaling

Previous work has shown that GIV mediates the activation of Akt downstream of various receptor tyrosine kinases (RTKs), including the Epidermal Growth Factor Receptor (EGFR), and other surface receptors via G-protein (i.e., Gβγ) dependent activation of PI3K (8, 15, 17, 23, 25, 33–35). We found that IGGi-11me reduced the phosphorylation of Akt at S473 (pAkt) upon EGF stimulation in two cell lines, MDA-MB-231 and HeLa, indicating reduced Akt activity (**Fig. 4A**). The lack of complete Akt inhibition is consistent with the known existence of GIV-independent mechanisms utilized by EGFR to activate PI3K-Akt signaling (36). In fact, the extent of IGGi-11me mediated inhibition of Akt was similar to that observed upon depletion of GIV in these cell lines (**Fig. 4B**). Moreover, IGGi-11me failed to further reduce Akt activation in GIV-depleted cells, indicating that it does not affect GIV-independent mechanisms of Akt activation downstream of EGFR (**Fig. 4B**). Also, IGGi-11me did not change the total amount of GIV or Gαi (**Fig. 4A, B**), supporting that its mechanism of action is the disruption of the interaction of the two proteins, rather than indirectly altering their abundance. These results show that IGGi-11me specifically inhibits GIV-dependent G-protein signaling in cancer cells.

**Figure 4.**
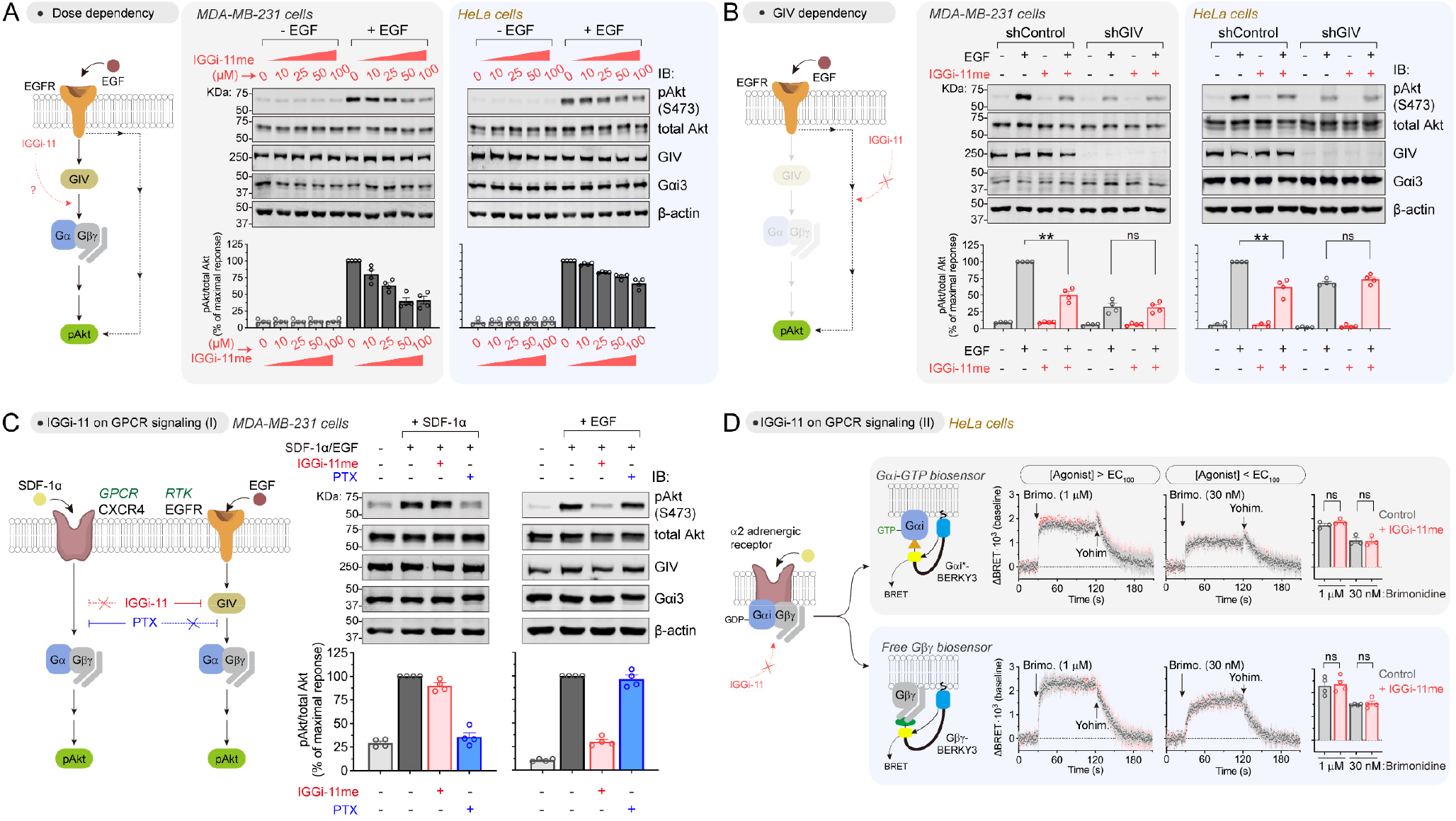
IGGi-11me specifically inhibits GIV-dependent G-protein cell signaling. **(A)** IGGi-11me inhibits EGF-stimulated Akt activation (phospho-serine 473, pAkt S473) in MDA-MB-231 and HeLa cells. Cells were preincubated with the indicated concentrations of IGGi-11me and stimulated with EGF (1.6 nM for MDA-MB-231 or 50 nM for HeLa) for 5 minutes before lysis and immunoblotting. **(B)** IGGi-11me (100 μM) does not inhibit EGF-stimulated Akt activation in GIV-depleted cells. GIV-depleted cells (shGIV) or control cells (shControl) were treated as in A. **(C)** IGGi-11me does not block Akt activation upon stimulation of the GPCR CXCR4. MDA-MB-231 cells were preincubated with IGGi-11me (100 μM) or pertussis toxin (PTX, 100 ng/ml) and stimulated with SDF-1α (100 ng/ml for 10 min) or EGF (1.6 nM for 5 min) before processing as in A. **(D)** IGGi-11me does not affect GPCR-mediated modulation of G-protein activity. HeLa cells expressing BRET biosensors for Gαi-GTP (Gαi*-BERKY3) or free Gβγ (Gβγ-BERKY3) were preincubated with IGGi-11me (100 μM) and sequentially treated with the α2 adrenergic agonist brimonidine and the antagonist yohimbine (25 μM) during real-time BRET measurements as indicated in the figure. All results are mean ± SEM (*N* ≥ 3). ** *P* < 0.01; ns, *P* > 0.05, ANOVA.

### IGGi-11me does not affect GIV-independent G-protein cell signaling

Although IGGi-11 does not interfere with GIV-independent mechanisms of G-protein regulation *in vitro* (**Fig. 3**, **Fig. S4-6**), confirmation that the same holds for IGGi-11 me in cells was warranted. First, we compared side by side the effect of IGGi-11me on GIV-dependent and GIV-independent G-protein signaling in the same cell line (MDA-MB-231) with the same readout (Akt activation). For GIV-dependent G-protein signaling, we stimulated cells with EGF as in **Fig. 4A-B**, whereas for GIV-independent G-protein signaling we stimulated cells with SDF-1α, an agonist for the endogenously expressed GPCR CXCR4 (**Fig. 4C**). We found that IGGi-11me inhibited Akt activation in response to EGF but not in response to SDF-1α (**Fig. 4C**), indicating that it does not interfere with GPCR-mediated G-protein signaling. In contrast, pertussis toxin (PTX), which precludes Gαi activation by GPCRs but not by GIV (37), efficiently blocked activation of Akt in response to SDF-1α but not to EGF (**Fig. 4C**). These results indicate that IGGi-11me specifically targets GIV-dependent G-protein signaling mechanisms in cells without interfering with canonical GPCR-mediated G-protein signaling. To further substantiate this point, we assessed the effect of IGGi-11me on GPCR signaling by using BRET-based biosensors that directly monitor the activation of endogenous G-proteins (31). More specifically, HeLa cells expressing biosensors for either Gαi-GTP or free Gβγ were treated with IGGi-11 me exactly under the same conditions that led to decreased GIV-dependent Akt activation after EGF stimulation (**Fig. 4A-B**). We found that G-protein responses elicited by stimulation of endogenous α2 adrenergic receptors with maximal (>EC100) or submaximal (<EC100) concentrations of a cognate agonist were unaltered by IGGi-11 me (**Fig. 4D**). Not only were the amplitudes and rates of the activation responses unchanged, but the rates of deactivation upon GPCR blockade with an antagonist also remained the same (**Fig. 4D**). These results show that IGGi-11me does not interfere with GIV-independent G-protein signaling like that elicited by GPCRs.

### IGGi-11me specifically inhibits GIV-dependent tumor cell migration

Previous evidence indicates that GIV is expressed at high levels in metastatic cancers, and that formation of the GIV-Gαi complex favors cell migration (15, 18–20). Consistent with some of these observations, we found that invasive breast cancer (BRCA) cell lines prone to metastasis expressed higher levels of GIV (GIV^High^) than non-invasive breast cancer cell lines (GIV^Low^) (**Fig. 5A**). IGGi-11me was approximately four times more potent inhibiting the migration of MDA-MB-231 cells (GIV^High^) than of MCF-7 cells (GIV^Low^) (**Fig. 5B**). This difference in IGGi-11me sensitivity could not be attributed to differences in Gαi protein abundance because they were present in similar amounts in both cell lines (**Fig. 5B**). While we could not test the effect of IGGi-11me on the GIV^Low^ cell lines T47D and MDA-MB-453 because they lacked measurable migration, we found that IGGi-11me inhibited cell migration in the GIV^High^ cell lines BT-549 and Hs578T with a potency similar to that seen for MDA-MB-231 cells (**Fig. S8A**). These results established a correlation between GIV expression (and presumably the formation of a GIV-Gαi complex) and sensitivity to IGGi-11me. To further substantiate that IGGi-11me specifically inhibits GIV-dependent tumor cell migration, we tested its effect on GIV-depleted MDA-MB-231 cells. We found that, compared to control cells, IGGi-11 me had no effect on MDA-MB-231 cell migration upon GIV depletion (**Fig. 5C**). Similar observations were made with HeLa cells (**Fig. 5D**). GIV-depleted cells contained amounts of Gαi proteins similar to those in their corresponding control cells (**Fig. 5C, D**), further supporting that the inhibition of cell migration exerted by IGGi-11me is GIV-dependent. Furthermore, the inhibition of cell migration by IGGi-11me was not a consequence of reduced cell viability, as the latter was not affected by the compound in any of cell lines investigated (**Fig. 5E**, **Fig. S8B**). These findings indicate that IGGi-11me specifically blocks GIV-dependent tumor cell migration, implying that the disruption of the GIV-Gαi complex hinders the pro-invasive features of GIV^High^ cancer cells.

**Figure 5.**
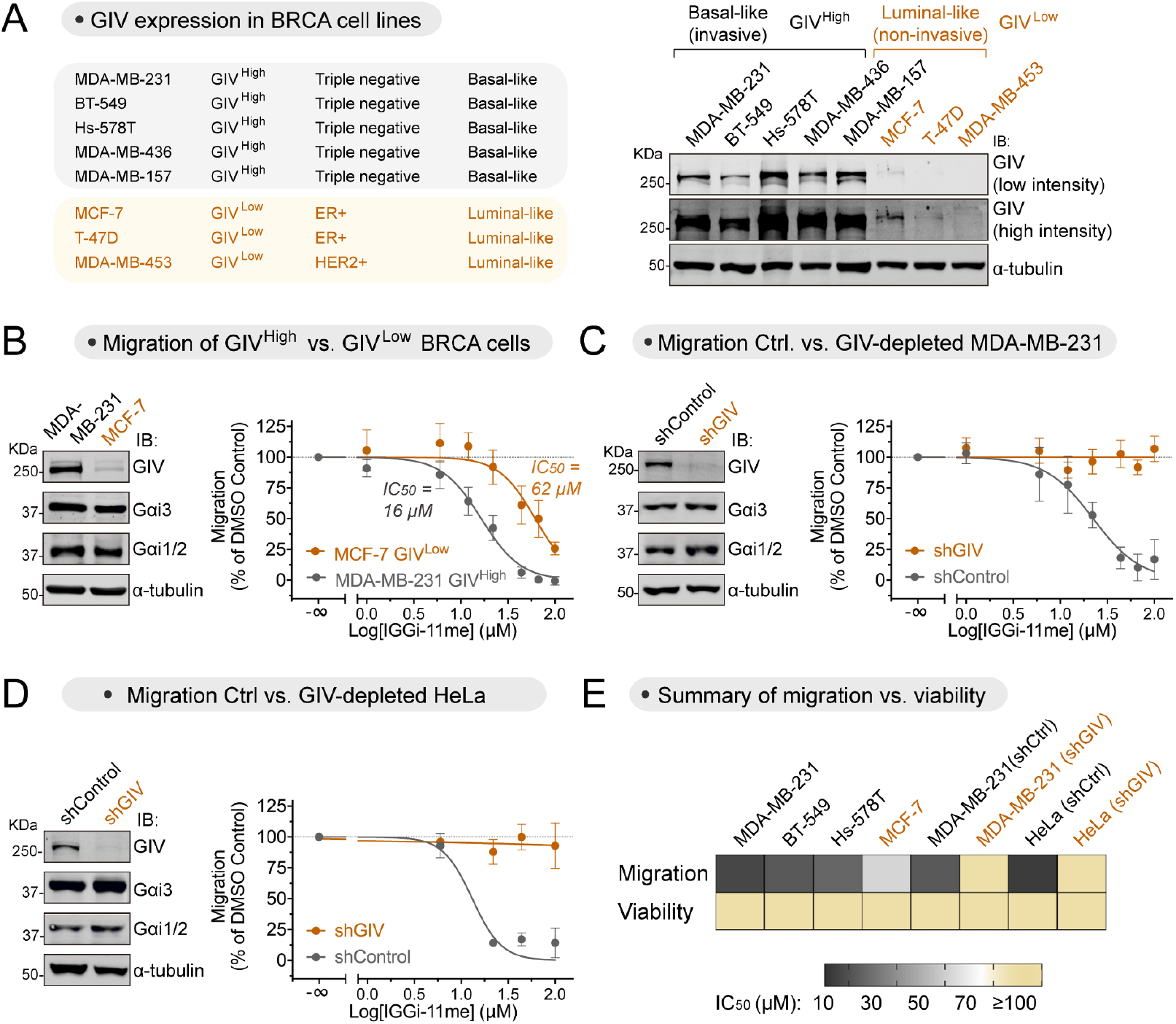
IGGi-11me blocks GIV-dependent tumor cell migration. **(A)** Basal-like invasive breast cancer (BRCA) cell lines express higher amounts of GIV (GIV^High^) than luminal-like non-invasive BRCA cell lines (GIV^Low^) as determined by immunoblotting. **(B)** IGGi-11me inhibits cell migration more potently in MDA-MB-231 cells (GIV^High^) than in MCF-7 cells (GIV^Low^). Chemotactic cell migration towards fetal bovine serum was determined in the presence of the indicated concentrations of IGGi-11me using a Boyden-chamber assay. **(C, D)** IGGi-11 me mediated inhibition of tumor cell migration is lost upon depletion of GIV from MDA-MB-231 (C) or HeLa (D) cells. GIV-depleted cells (shGIV) or control cells (shControl) were processed as described in B. **(E)** IGGi-11me impairs tumor cell migration without affecting cell viability. Heatmap comparing the half-maximal inhibitory concentration (IC_50_) of IGGi-11me on cell migration or viability of the indicated cell lines. IC_50_ values were determined from results shown in this figure or in **Fig. S8**. Cell viability was determined upon incubation with IGGi-11me for 24 hours, which is longer than the times cells were exposed to the compound in cell migration assays. All results are mean ± SEM (*N* ≥ 3).

### IGGi-11me inhibits tumor growth and metastatic invasion in mice

Although loss of GIV does not affect the growth of tumor cells on plastic dishes, it hinders growth in three-dimensional Matrigel cultures (17), which account for tumor cell interactions with the extracellular matrix and recapitulate many of the behavioral features of cancer cells in tumors *in situ* (38). We found that IGGi-11me mimicked previous observations (17) upon loss of GIV in Matrigel cultures— i.e., MDA-MB-231 became smaller and more organized acinar structures than control cells, resulting in an overall reduction of cell growth (**Fig. 6A, B**). In contrast, IGGi-11me did not affect the growth of non-transformed MCF-10A breast cells in Matrigel cultures (**Fig. 6B**). Consistent with these observations *in vitro*, IGGi-11me also reduced the ability of MDA-MB-231 cells to form tumors when implanted subcutaneously as xenografts in mice (**Fig. 6C**). Because we could not observe metastatic invasion of the lungs upon subcutaneous tumor implantation, we assessed the effect of IGGi-11me on MDA-MB-231 cell injection through the tail vein, an established experimental paradigm of metastatic colonization of the lungs (39). We found that IGGi-11me reduced the ability of MDA-MB-231 cells to appear in the lungs weeks after tail vein injection (**Fig. 6D**). Together, these results indicate that disruption of the GIV-Gαi interaction by IGGi-11me prevents tumor cell traits associated with cancer progression.

**Figure 6.**
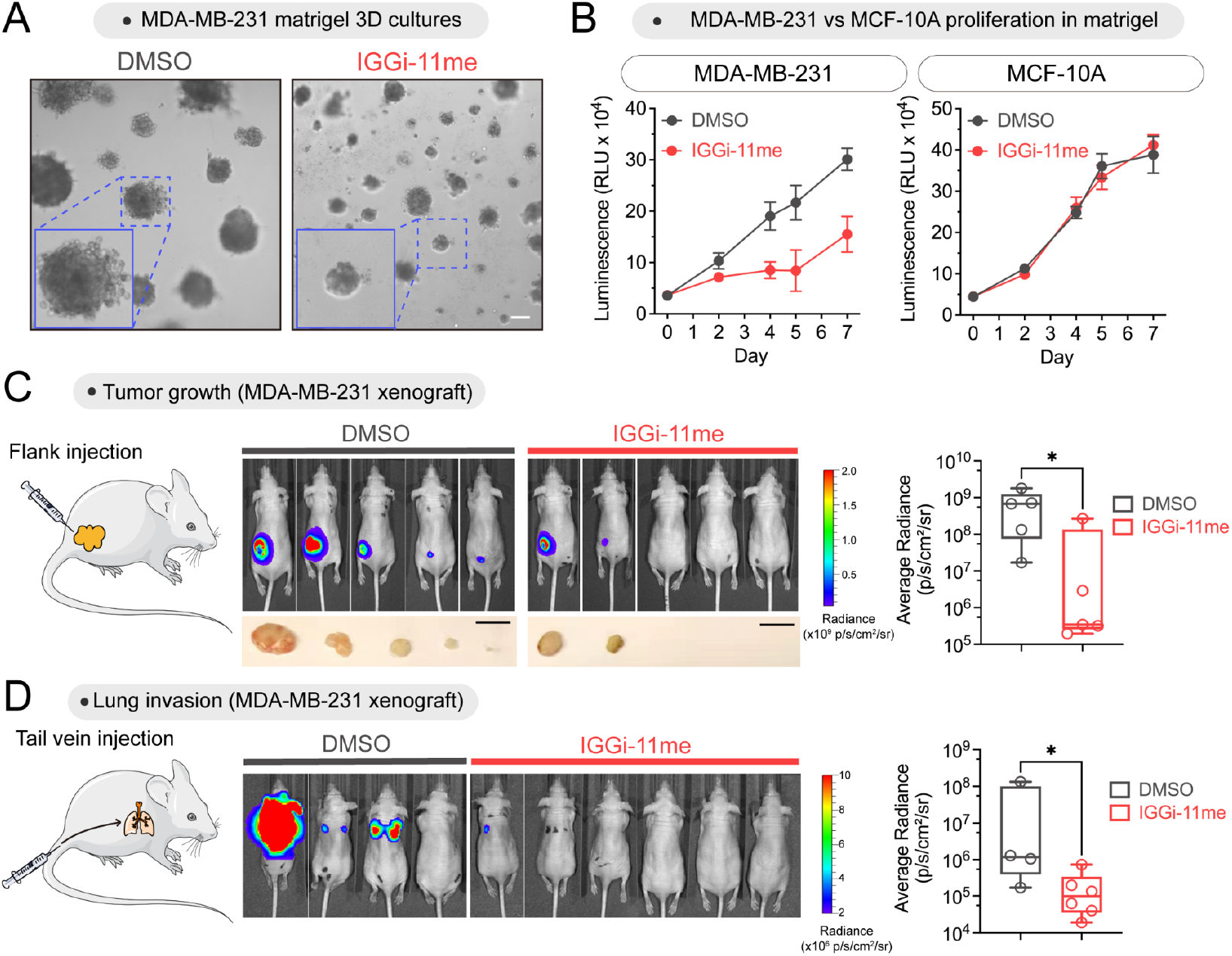
Inhibition of tumor growth and metastatic invasion in mice upon IGGi-11me treatment. **(A, B)** IGGi-11me inhibits growth of MDA-MB-231 breast cancer invasive cells, but not of non-transformed MCF-10A cells, on Matrigel. IGGI-11 me (100 μM) or DMSO was used to treat cells at the onset of the culture period for 2 days and then removed for the remaining duration of the experiment. (A) displays representative images of acini at 7 days (scale bar = 100 μm), and viability in (B) is expressed as mean ± SEM (*N* ≥ 3). **(C, D)** IGGi-11 me impairs MDA-MB-231 cell tumor growth (C) or lung invasion (D) in NCr nu/nu athymic nude mice. Female nude mice were injected subcutaneously (C) or through the tail vein (D) with luciferase-expressing MDA-MB-231 cells treated with IGGi-11me or DMSO, and imaged 8 weeks later upon luciferin administration (*N* = 4-6 per group). Box plots on the left display the quantification of luminescence (median, min/max). *\*P* < 0.05, Mann-Whitney U test. In (C), tumors were photographed post-mortem (scale bar = 1 cm).

## Discussion

In this work we identify and characterize a chemical probe of broad utility for dissecting atypical mechanisms of cellular communication mediated by G-proteins with important biomedical implications not only for cancer, but also for fibrosis, and male fertility, among other maladies (8, 9, 17, 18, 20, 40, 41). Because its chemical scaffold is synthetically tractable, IGGi-11 may further serve as a lead compound to develop analogs with improved potency and pharmacokinetic properties that could have therapeutic value. From a broader perspective, this work provides the proof of principle for a modality of pharmacological targeting in heterotrimeric G-protein signaling that deviates from the widespread focus on GPCRs or the direct ablation of G-protein activity *en toto*. This modality consists of targeting G-proteins to selectively disrupt specific mechanisms by which they are regulated. IGGi-11 disrupts Gαi binding to GIV but not to many of its other binding partners, despite them physically engaging the same region of Gαi as GIV. This region includes the SwII, which is dynamic and adopts different conformations depending on the protein partner bound to Gαi. Although it is tempting to speculate that the selectivity of IGGi-11 may arise from its relative ability to interact with these different conformations, the structural basis for the action of IGGi-11 remains to be fully elucidated. The targeting modality described here follows the path opened by recent advances on small molecule inhibitors for another GTPase, KRas, in reshaping the traditional definition of what constitutes a druggable target (42, 43).

## MATERIALS and METHODS

Chemical compounds of interest were purchased from reliable vendors or synthesized in-house, and tested in *in vitro* assays, including nuclear magnetic resonance (NMR), bioluminescence resonance energy transfer (BRET) assays or different protein-protein binding experiments following previously established procedures that are described in details in Supplementary Information. Cell-based experiments to assess the efficacy and specificity of compounds were also carried out using previously established procedures and/or cell lines, including cell migration assays using modified Boyden chambers, immunoblotting and signaling assays, all of which are described in detail in Supplementary Information along with the animal experiments measuring xenograft tumor growth by luminescence bioimaging.

## ACKNOWLEDGEMENTS

This work was supported by NIH grant R01GM130120 and the Karin Grunebaum Cancer Research Foundation (to MG-M). JZ is supported by a Dahod International Scholar Award, and AL is supported a F31 Ruth L. Kirschstein NRSA Predoctoral Fellowship (F31NS115318). FJB is supported by Spanish Government grant PID2020-113225GB-I00. MF-G is supported by Spanish Government fellowship PRE-2018-085788. We thank the ICCB-Longwood Screening Facility at Harvard Medical School, S. Whelan (Boston University), F. Seta (Boston University), N. Ganem (Boston University), and J.B. Blanco-Canosa (Institute for Advanced Chemistry of Catalonia) for access to instrumentation and reagents. We thank N. Merino (CIC bioGUNE, Spain) for help with the purification of Gαi3 protein used in NMR studies, M. Rico (CIC bioGUNE, Spain) for access to NMR spectrometers, and A. González-Magaña (CIC bioGUNE, Spain) for preliminary ITC experiments. We thank the following investigators for providing DNA plasmids: K. Martemyanov (The Scripps Research Institute, Jupiter, FL), N. Lambert (Augusta University, Augusta, GA), P. Wedegaertner (Thomas Jefferson University, Philadelphia, PA), J. Blumer (Medical University of South Carolina, Charleston, SC), J. Sondek (University of North Carolina, Chapel Hill, NC), N. Artemyev (University of Iowa) C. Dressauer (University of Texas Health Science Center at Houston, TX), and M. Linder (Cornell University).

## Author Contributions

J.Z., V.D., M.F.G, S.D., J.-C.P., A.L., and Q.C. conducted experiments. J.Z., V.D., A.B., F.J.B., and M.G-M. designed experiments. J.Z., V.D., M.F.G, S.D., J.-C.P., A.L., A.I.dO, and F.J.B. analyzed data. M.G.-M. wrote the manuscript with input from all authors. M.G.-M. conceived and supervised the project.

## Competing interests

M.G.-M. is listed as an inventor in a provisional patent filed by Boston University related to the content of this manuscript. The rest of the authors declare no competing interests.

## Data and materials availability

All data are available in the main text or the supplementary materials.

## Notes

### Competing Interest Statement

Boston University has filed a provisional patent application related to the content of this manuscript in which Mikel Garcia-Marcos is listed as an inventor

