## Supplemental information for "Small-molecule targeting of GPCR-independent non-canonical G protein signaling inhibits cancer progression"

**Supplementary information includes 8 figures and 1 table**

Materials and Methods

Figs. S1 to S8

Tables S1

Supplemental references

#### MATERIALS and METHODS

##### Synthesis of chemical compounds

IGGi-11 (4'-((9H-fluorene-2,7-disulfonyl)bis(methylazanediyl))dibutyric acid) was purchased from Chembridge or Sigma (R693073), or synthesized as follows. Synthesis of IGGi-11me from IGGi-11 is also described below.

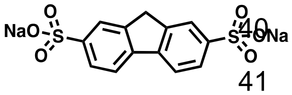

Chlorosulfonic acid (1 ml, 20 mmol, 3 equiv) was added dropwise at 0 °C to a stirred solution of fluorene (1 g, 6 mmol, 1 equiv) in acetic acid (10 ml, 0.6 M). The reaction mixture was refluxed for 2 h, cooled and poured into a saturated aqueous solution of NaCl (10 ml) containing NaOH (600 mg, 2.5 equiv) to obtain a yellow precipitate (1.9 g, 90%). The precipitate was washed three times with a saturated solution of NaCl, filtered and dried overnight at 60 °C to give disodium 9H-fluorene-2,7-disulfonate as an off-white solid (1.9 g, 90%).

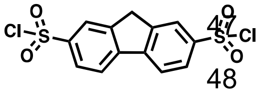

Sodium 9H-fluorene-2,7-disulfonate (2g, 5 mmol, 1 equiv), $PCl_5$ (3g, 20 mmol, 3 equiv) and $POCl_3$ (6 mL, 60 mmol, 12 equiv) were heated to reflux for 16 h. $POCl_3$ was removed by distillation before water was added slowly and the mixture was sonicated to break up the aggregates. The solid obtained was filtered to obtain 9H-fluorene-2,7-disulfonyl dichloride as a brown solid (1.87g, 100%).

**$^1$H NMR (500 MHz, $CDCl_3$)** δ 8.31 (s, 2H), 8.18 (d, *J* = 7.9 Hz, 2H), 8.11 (d, *J* = 8.2 Hz, 2H), 4.21 (s, 2H).

**$^{13}$C NMR (126 MHz, $CDCl_3$)** δ 145.62, 145.41, 144.23, 126.77, 124.11, 122.26, 37.30.

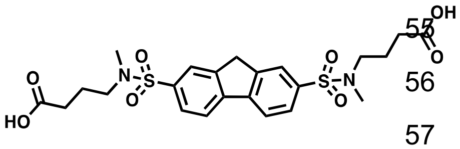

4-(Methylamino)butanoic acid (19 mg, 0.17 mmol, 2 equiv) and DIPEA (43 µl, 0.25 mmol, 3 equiv) were added to a 1-dram vial containing 9H-fluorene-2,7-disulfonyl dichloride (30 mg, 0.08 mmol, 1 equiv) in DMA (1 mL). The mixture was stirred at 23 °C for 12 h until the consumption of the starting material was observed by TLC (DCM: MeOH, 97:3). The mixture was then quenched with 1M HCl (1.2 ml) and was extracted three times with DCM and washed three times with brine. The crude was dried over $Na_2SO_4$ and the solvent was evaporated *in vacuo* and purified by column chromatography (DCM: MeOH, 97:3) to afford a white solid (35 mg, 81%, >95% purity).

**$^1$H NMR (500 MHz, $CD_3OD$)** δ 8.15 (d, *J* = 8.1 Hz, 2H), 8.07 (s, 2H), 7.88 (dd, *J* = 8.1, 1.7 Hz, 2H), 4.16 (s, 2H), 3.10 (t, *J* = 6.9 Hz, 4H), 2.77 (s, 6H), 2.37 (t, *J* = 7.2 Hz, 4H), 1.83 (p, *J* = 7.1 Hz, 4H).

**$^{13}$C NMR (126 MHz, $(CD_3)_2SO$)** δ 173.22, 145.27, 143.94, 137.22, 126.57, 124.40, 121.33, 49.41, 36.91, 34.34, 29.76, 22.64.

**HR-MS (*m/z*):** $[C_{23}H_{28}N_2O_8S_2+H]^+$ calculated: 525.1365; found: 525.1355 (+1.9043 ppm).

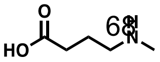

4-(Methylamino)butanoic acid used in the reaction above was synthesized as follows: hydrochloric acid (7 ml, 7.2 M) was added to 1-methylpyrrolidin-2-one (5 g, 50 mmol, 1 equiv) and the mixture was heated to reflux for 16 h. The HCl was evaporated in Genevac and the crude material was purified by column chromatography (DCM: 10% $NH_4OH$/MeOH, 90:10) to afford the final product as an off-white solid (4 g, 70%).

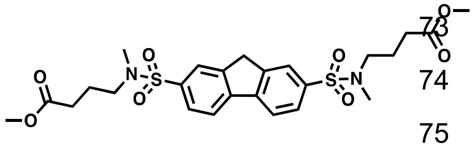

Thionyl chloride (5.7 µl, 79 µmol, 2.06 equiv) was added to a solution of 4,4'-((9*H*-fluorene-2,7-disulfonyl)bis(methylazanediyl))dibutyric acid (20 mg, 38 µmol, 1 equiv) in MeOH (1 ml) at 0 °C, and the solution stirred for 1.5 h. The solvent was evaporated *in vacuo* to afford the final product **IGG-11me** as a white solid with no further purification (21.5 mg, quantitative, >95% purity).

**$^1$H NMR (400 MHz, $CDCl_3$)** δ 8.00 (s, 2H), 7.96 (d, *J* = 8.1 Hz, 2H), 7.85 (d, *J* = 8.0 Hz, 2H), 4.07 (s, 2H), 3.68 (s, 6H), 3.10 (t, *J* = 6.8 Hz, 4H), 2.77 (s, 6H), 2.44 (t, *J* = 7.2 Hz, 4H), 1.88 (p, *J* = 7.1 Hz, 4H).

**$^{13}$C NMR (101 MHz, $CDCl_3$)** δ 126.74, 124.29, 121.20, 53.77, 51.69, 49.40, 42.04, 34.76, 30.61, 22.69, 18.63, 17.37, 12.02.

**HR-MS (*m/z*):** $[C_{25}H_{32}N_2O_8S_2+H]^+$ calculated: 553.1678; found: 553.1688 (+1.8078 ppm).

All $^1$H NMR spectra were recorded at 400 or 500 MHz at ambient temperature with $CDCl_3$, $CD_3OD$, DMSO ($d_6$) or $D_2O$ as the solvent. Chemical shifts are recorded in parts per million (ppm) relative to $CDCl_3$ ($^1$H, δ 7.26; $^{13}$C, δ 77.1), $CD_3OD$ ($^1$H, δ 3.31; $^{13}$C, δ 49.0), $(CD_3)_2SO$ ($^1$H, δ 2.50; $^{13}$C, δ 39.5) or $D_2O$ ($^1$H, δ4.79). Analytical LC-MS was performed on a Waters Acquity UPLC (Ultra Performance Liquid Chromatography (Waters MassLynx Version 4.1) with a Binary solvent manager, SQ mass spectrometer, Water 2996 PDA (PhotoDiode Array) detector, and ELSD (Evaporative Light Scattering Detector).

**Reconstitution of compound solutions**

Powder stocks of IGGi-11 were resuspended in DMSO at a final concentration of 100 mM and directly diluted in aqueous solutions up to a 1 mM concentration for experiments. Powder stocks of IGGi-11me were resuspended in DMSO at a final concentration of 40 mM and diluted in aqueous solutions for biological experiments as described next. One-thousand and five hundred µl of cell culture media were added to a 15 ml conical tube with 3.75 µl of 40 mM IGGi-11me at the bottom, and pipetted up and down 8-10 times for mixing. This solution was incubated at room temperature for 10 min in a sonicated bath (EMERSON, Branson Bransonic® CPXH Digital Bath 1800). This solution (100 µM) was used to make serial dilutions as needed. Cell media was replaced by IGGi-11me media for compound treatments.

The sources and structures of all other "IGGi" compounds (69 including IGGi-11) are presented in **Table S1**. All these compounds were resuspended in DMSO at a final concentration of 40 mM.

**Plasmids**

*E. coli* expression plasmids encoding His-tagged rat Gαi3 (rat His-Gαi3; pET28b-rGαi3), His-tagged human Gαi3 (human His-Gαi3; pET24d-hGαi3), His-tagged rat Gαi2 (rat His-Gαi2; pET28b-rGαi2), His-tagged human Gαi1 (human His-Gαi1; pPro-Gαi1), GST-tagged rat Gαi3 (GST-Gαi3; pGEX-4T-1-GST-Gαi3), His-tagged human GIV-CT (His-GIV-CT (aa 1660-1870); pET28b-hGIV (1660-1870)), and GST-tagged human GST-GIV (1671-1755; pGEX-4T-1 hGIV (1671-1755)) have been described previously (1-7). Plasmids encoding His-tagged RGS4 (His-RGS4, pLIC-His-RGS4), and GST-tagged RGS4 (GST-RGS4, pLIC-GST-RGS4), were generated using a ligation independent cloning (LIC) procedure (8). This procedure was used to insert the sequence encoding human RGS4 into pLIC-His or pLIC-GST vectors kindly provided by J. Sondek (UNC-Chapel Hill, NC) (9). The plasmid encoding His-tagged bovine Gαs (pHis6-Gαs) was kindly provided by N. Artemyev (University of Iowa). The plasmid for producing myristoylated rat Gαi1 used in adenylyl cyclase experiments was generated by amplifying the sequence of Gαi1 with an internal His-tag in the b/c loop from a pQE-Gαi1H6 plasmid provided by C. Dressauer (University of Texas Health Science Center at Houston, TX) and inserting it in the NdeI/BglII sites of pLIC-His using Gibson assembly (pLIC- Gαi1-int6xHis). The pbb131 plasmid encoding N-myristoyltransferase (NMT) (pbb131-NMT) was a gift from M. Linder (Cornell University) (10). All point mutations were generated using QuikChange II (Agilent, #200523). Mammalian expression plasmids encoding the BRET acceptor Venus-tagged Gβγ (pcDNA3.1-Venus[1-155]-$G\gamma_2$[VN-$G\gamma_2$] and pcDNA3.1-Venus[155-239]-$G\beta_1$[VC-$G\beta_1$]) or untagged Gβγ (human $G\beta_1$; pcDNA3.1-$G\beta_1$, and human $G\gamma_2$; pcDNA3.1-$G\gamma_2$) were kindly provided by N. Lambert (Augusta University, Augusta, GA) (11, 12), and the plasmid encoding bovine GRK3ct (aa 495–688) fused to nanoluciferase and a membrane anchoring sequence, “mas” (mas-GRRK3ct-Nluc; pcDNA3.1-masGRK3ct-NanoLuc) was a kind gift from K. Martemyanov (Scripps Research Institute, Jupiter, FL) (13). The plasmids encoding untagged rat Gαi3 (pcDNA3-Gαi3) and untagged rat Gαo (pcDNA3-Gαo) have been described previously (14, 15). The plasmid encoding Gαq-HA (mouse, pcDNA3-Gαq-HA, internally tagged) was kindly provided by P. Wedegaertner (Thomas Jefferson University) (16). The plasmids encoding human Gα13 (pcDNA3.1-Gα13 internally EE-tagged; cat#GNA130EI00) or human M3R (pcDNA3.1-3xHA-M3R; cat#MAR030TN00) were obtained from the cDNA Resource Center at Bloomsburg University. The plasmid encoding rat $\alpha2_A$-AR (pcDNA3-$\alpha2_{A/D}$-AR)(17) was provided by J. Blumer (Medical University of South Carolina, SC), and the plasmid for PAR1 (18) was obtained from Addgene (pBJ-FLAG-hPAR1 #53226). The plasmid encoding rat Gαi3 with citrine variant of YFP inserted in the αb/αc loop of Gαi3 (pcDNA3.1-Gαi3-YFP (b/c loop) has been described previously (19). The plasmid encoding human adenylyl cyclase 5 (pcDNA3.1-YFP-hAC5) was a kind gift from C. Dessauer (University of Texas Health Science Center at Houston, TX) (20). The plasmids encoding GIV or control shRNA sequences (pLKO.1-puro-GIV shRNA2, AAGAAGGCTTAGGCAGGAATT; pLKO.1-puro-scr, GGATTGAGATCAGAAGATAGC) have been described previously (*25*). The lentiviral plasmid encoding myc-tagged firefly luciferase (pLVX- fluc2-myc-IRES-Hyg) was generated in two steps. First, firefly luciferase (fluc2) was amplified by PCR from pGL4.33 (Promega, cat# E1340) and inserted into the NheI/KpnI of pcDNA3.1(+) to generate pcDNA3.1-fluc2-myc. Then, the fluc2-myc cassette was PCR amplified and inserted into the XhoI/BamHI sites of pLVX-IRES-Hyg (Clontech, cat# 632185) by Gibson assembly to generate pLVX-fluc2-myc IRES-Hyg. Lentiviral packaging plasmids were pSPAX2 (Addgene #12260) and pMD2.G (Addgene #12259).

**Protein purification and peptide synthesis**

Purification of rat His-Gαi3, human His-Gαi3, rat His-Gαi2, human His-Gαi1, human His-GIV-CT (aa 1660-1870), human His-RGS4, rat GST-Gαi3, human GST-GIV (aa 1671-1755), or human GST-RGS4 proteins was carried out as described previously (2, 3) with minor modifications. Briefly, protein expression was induced in BL21(DE3) *E. coli* cells transformed with the appropriate plasmids by overnight incubation with 1 mM isopropyl-β-D-1-thio-galactopyranoside (IPTG) at 23 °C, except for rat His-Gαi3 used in the high-throughput fluorescence polarization experiments, which was induced overnight at 23 °C using the Studier's autoinduction method (21). Bacteria pelleted from 1 liter of culture were resuspended in 25 mL of lysis buffer (50 mM $NaH_2PO_4$, 300 mM NaCl, 10 mM imidazole, 1% (v:v) Triton X-100, 1 μM Leupeptin, 2.5 μM Pepstatin, 0.2 μM Aprotinin, 1 mM PMSF, pH 7.4). For G-protein preparation, the buffer was supplemented with 25 μM GDP and 5 mM $MgCl_2$. After sonication (four cycles, with pulses lasting 20 s/cycle, and with 1 min interval between cycles to prevent heating), lysates were centrifuged at 12,000 × g for 20 min at 4 °C. Solubilized proteins were affinity purified on either HisPur Cobalt resin (ThermoFisher cat#89964) for His-tagged proteins or Glutathione Agarose resin (Pierce 16100) for GST-tagged proteins, and eluted with lysis buffer supplemented with 250 mM imidazole or with 50 mM Tris-HCl, 100 mM NaCl, 30 mM reduced glutathione (pH 8), respectively. Eluted proteins were dialyzed overnight at 4°C against PBS, except for G-proteins, which were buffer exchanged to 20 mM Tris-HCl, 20 mM NaCl, 1 mM $MgCl_2$, 1 mM DTT, 10 μM GDP, 5% (v/v) glycerol (pH 7.4) on an FPLC system using a HiTrap Desalting column (GE Healthcare cat# 17-1408-01). For human His-Gαi3 used in isothermal titration calorimetry experiments, buffer exchange was to 10 mM HEPES, 10 mM $MgCl_2$, 1 mM TCEP, 300 μM GDP (pH 7) using the same desalting column. His-tag-cleaved human Gαi3 used for nuclear magnetic resonance experiments (2), bovine His-Gαs (22), and myristoylated rat Gαi1 with an internal His-tag (23) were purified exactly as in the indicated references. All protein preparations were aliquoted and stored at -80 °C. Fluorescently (FITC) labeled peptides corresponding to human GIV (aa 1671-1701, KTGSPGSEVVTLQQFLEESNKLTSVQIKSSS), RGS12 GoLoco motif (aa 1185-1221, DEAEEFFELISKAQSNRADDQRGLLRKEDLVLPEFLR, R12 GL), or the synthetic sequence KB-1753 (SSRGYYHGIWVGEEGRLSR) were synthesized and purified exactly as described previously (2, 3, 24), dissolved in DMSO, and stored in aliquots at -80 °C.

**High-throughput screen by fluorescence polarization**

This screen was conducted at the ICCB-Longwood Screening Facility at Harvard Medical School with a collection of ~200,000 compounds from commercial and academic sources. The principle of the assay is based on monitoring binding of a GIV-derived peptide to Gαi3, which has been previously shown to recapitulate the properties the native GIV-Gαi3 interaction (2, 3). Experimental wells of 384-well assay plates (black ProxiPlate F-Plus, Perkin Elmer cat# 6008269) were pre-filled with FITC-GIV (aa 1671-1701) peptide in a 10 μl volume of assay buffer (50 mM Tris, 100 mM NaCl, 10 mM $MgCl_2$, 5 mM EDTA, 0.4% (v:v) NP-40, 30 μM GDP, 1 mM DTT, pH 7.4) using a Multidrop Combi liquid handler (Thermo). Plates were centrifuged at 1000xg for 2 minutes to eliminate bubbling on the surface, and 100 nl of experimental compound (in 100% DMSO) was pin-transferred into individual wells in two replicate plates using a D-TRAN XM3106-31 PN robot (Seiko). After pin-transfer, 5 μl of rat His-Gαi3 was added to each well using a Multidrop Combi liquid handler (Thermo), and assay plates were shaken at low speed for 5 seconds. Plates were centrifuged at 1000xg for 2 minutes and shaken for 10 seconds prior to an incubation of 90 minutes at room temperature. The final concentrations of assay components were 25 nM for FITC-GIV and 1 μM for His-Gαi3. The typical final compound concentration was approximately 30 μg/ml, with a final concentration of DMSO per well of 0.67% (v:v). Fluorescence polarization (FP) signal was measured with an EnVision Multilabel 2103 plate reader (PerkinElmer) using a D505fp/D535 dual mirror (Ex 480 nm / Em 535 nm filters, P and S channels). Each plate contained 16 negative control wells containing only 0.67% DMSO without test compound and 16 positive control wells containing 30 μM $AlCl_3$ and 10 mM NaF (to generate the $AlF_4^-$ species that completely disrupts the GIV-Gαi3 interaction in this assay format (3)). Negative and positive controls were used to normalize the FP signals to 100% and 0% GIV-Gαi3 binding, respectively. Compounds reducing binding 15% or more were considered hits (~580 compounds). Hits were re-tested in this assay format using the same procedure and in an AlphaScreen® assay described below, substituting a D300e liquid dispenser (Hewlett Packard) for compound addition (200 nl). Results of the screen were deposited in PubChem (AID: 1224905).

**AlphaScreen® assay**

As an approach orthogonal to monitoring the GIV-Gαi interaction by fluorescence polarization, we implemented a previously established AlphaScreen® assay. This chemiluminescent assay operates at wavelengths different from those used in the fluorescence polarization assay to monitor the association between Gαi3 and a fragment of GIV that recapitulates the binding properties of the full-length protein (2, 3, 25). For AlphaScreen® experiments in high-throughput screening format, 200 nl of experimental compounds (“cherry-picked” from the original libraries) diluted in DMSO were added to individual wells of 384-well plates (white ProxiPlate F-Plus, Perkin Elmer cat#6008280) containing 5 μl of His-GIV-CT (aa 1660-1870) diluted in assay buffer (50 mM Tris, 100 mM NaCl, 5 mM $MgCl_2$, 0.4% (v;v) NP-40, 50 μM GDP, pH 7.4). Five μL of GST-Gαi3 diluted in assay buffer were added to each well, and plates were centrifuged at 1000xg for 2 minutes, shaken for 10 seconds, and incubated for 90 minutes at room temperature to allow for GIV-Gαi3 complex formation. Next, 5 μL of a suspension of AlphaScreen Nickel-chelate donor beads (Perkin Elmer, cat#AS101) and AlphaLISA Glutathione acceptor beads (Perkin Elmer, Cat#AL109) diluted in assay buffer were added to each well. Multidrop Combi liquid handlers (Thermo) were used for dispensing all buffers and reagents, except for compounds, which were delivered using a D300e liquid dispenser (Hewlett Packard). Each compound was tested in triplicate, and the final concentrations of components were as follows: 50-100 μM for compounds (fixed 200 nl volume of library stocks), 75 nM each of His-GIV-CT and GST-Gαi3, 10 μg/ml for AlphaScreen Nickel-chelate donor beads, and 5 μg/ml for AlphaLISA Glutathione acceptor beads. The final concentration of DMSO was 1% (v:v). Controls included in each run consisted of no compound, DMSO-only negative controls, and $AlF_4^-$ added with His-GIV-CT to disrupt the GIV-Gαi3 interaction (3) as positive controls (50 μM $AlCl_3$, 10 mM NaF final concentrations). Signals were read with an EnVision Multilabel 2103 plate reader (PerkinElmer) using the AlphaScreen®-rated D640as mirror and M570w emission filter (570 ± 50 nm), and normalized to negative (100%) and positive (0%) controls. Compounds reducing the AlphaScreen® signal 35% or more compared to negative controls (DMSO) were considered positive hits. Re-testing of freshly purchased compounds was carried out as above but in the presence of different doses of compound (0.05-100 μM).

**Triage of hit compounds**

Compounds that were deemed hits from both the fluorescence polarization and the AlphaScreen® assays were manually evaluated for known pan assay interference (PAINS) moieties and other electrophilic properties that would result in undesired promiscuity. We further excluded compounds with more stringent criteria after applying computational filters to screen for additional PAINS and problematic chemical groups (26, 27).

**Cell culture and establishment of cell lines**

MDA-MB-231 (ATCC #HTB-26), MCF-7 (ATCC #HTB-22), Hs-578T (ATCC #HTB-126), BT-549 (ATCC #HTB-122), MDA-MB-436 (ATCC #HTB-130), MDA-MB-157 (ATCC #HTB-24), T47-D (ATCC #HTB-133), MDA-MB-453 (ATCC #HTB-131), HeLa (ATCC #CCL-2), and HEK293T (ATCC #CRL3216) cells were maintained at 37 °C with 5% $CO_2$ in DMEM (Gibco cat#11-965-118) supplemented with 10% (v:v) fetal bovine serum (FBS), 100 U/ml penicillin, 100 μg/ml streptomycin, and 2 mM L-glutamine. The medium for some of these cell lines was supplemented as follows: Hs-578T, 0.01 mg/ml insulin; BT-549, 0.002 mg/ml insulin; MDA-MB-436, 0.002 mg/ml insulin; T47-D, 0.002 mg/ml insulin. MCF-10A cells (ATCC #CRL-10317) were maintained in MEGM (Lonza cat#CC3150) supplemented with components of the SingleQuot supplement kit (Lonza cat#C4136) and 100 ng/mL cholera toxin (List Biological Laboratories #100B). The FBS used for all cell lines was from Gibco (Cat# 26140-079), except for HEK293T cells, which were grown in FBS from

HyClone (Cat# SH30072.03). MDA-MB-231 stably expressing firefly luciferase were generated by lentiviral transduction followed by antibiotic selection.

Lentiviral particles were produced by transfection of HEK293T cells using polyethylenimine (PEI; Polysciences, Inc; #23966, 1 mg/ml solution reconstituted in water). Four hundred thousand cells were seeded per well of a 6-well plate and cotransfected the next day with lentiviral plasmid of interest (1.8 µg) and packaging plasmids psPAX2 (1.2 µg) and pMD2.G (0.75 µg). Plasmid DNA was added to 200 µl of fresh DMEM without serum and mixed with 7.5 µl of PEI reagent by vortexing for 2 seconds. Tubes were incubated at room temperature for 15 min before adding to cells. Six hours after transfection, the media was changed to DMEM with 10% FBS. Lentivirus-containing media were collected 24 hr and 48 hr after transfection and combined together, centrifuged at 1500x*g* for 5 min, and filtered through a 0.45-µm surfactant-free cellulose acetate (SFCA) membrane (Corning, cat#431220). MDA-MB-231 cells were seeded on 6-well plates (200,000 cells per well). The day after seeding, cells were transduced by a 48 hr incubation with 2 ml of a 1:1 mix of lentivirus-containing supernatants described above mixed with fresh complete media and supplemented with 6 µg/ml of polybrene. Cells were transferred to a 10-cm plate and selection with 250 µg/ml hygromycin (GoldBiotechnnology, cat#H-270-5) started the day after. All surviving clones were pooled and maintained in the presence of 250 µg/ml hygromycin.

The generation of MDA-MB-231 and Hela cells stably expressing GIV shRNA or a control shRNA (has been described previously (28). These cells were maintained in their culture medium supplemented with 1 µg/ml puromycin (GoldBiotechnnology, cat#P-600-1). HeLa cells stably expressing the biosensors Gαi*-BERKY3 or Gβγ-BERKY3 have also been described previously (29), and were maintained in their culture medium supplemented with 100 µg/ml hygromycin.

**Tumor cell migration assay**

Cell migration was evaluated using a modified Boyden chamber assay. MDA-MB-231, MCF-7, Hs-578T, BT-549, or HeLa cells were grown to approximately 70% confluency in a 10 $cm^2$ culture plate prior to detachment by warm citric saline solution (135 mM KCl, 15 mM Sodium Citrate, pH 7.2) with a 10 minute incubation at 37 °C after three washes with citric saline. Detached cells were washed with 15 ml medium supplemented with 0.2% FBS three times by cycles of centrifugation (300xg, 3 minutes), aspiration and resuspension. Cells were seeded into each well of the top chamber of a 96-well 8 µm polyester transwell migration plate (Corning cat#3374) in a 40 µl volume of medium supplemented with 0.2% FBS. Forty µl of the same medium (0.2% FBS) containing compounds at 2X of the final concentration indicated in the figures or figure legends were added to the wells and incubated for 1 hour at room temperature. After this serum-starved preincubation with compound, 250 µl of medium supplemented with 10% FBS was added to the bottom chamber to create the migratory gradient. The concentration of compound in the bottom chamber was matched to that in the upper chamber, and the concentration of DMSO was maintained constant at 0.5%

(v:v) across all conditions. Medium without FBS in the bottom chamber was used a negative control for migration. The initial number of cells seeded per well and the time of incubation at 37 °C in 5% $CO_2$ to allow the migration of cells was as follows: MDA-MB-231, 25,000 cells for 6 hours; MCF-7, 60,000 cells for 18 hours; Hs-578T, 25,000 cells for 16 hours; BT-549, 25,000 cells for 16 hours; HeLa, 25,000 cells for 16 hours. Following the migration period, the top chamber was cleared of non-migratory cells using a cotton swab and medium was aspirated with gentle vacuum. The top chamber was then moved to a receiver plate containing warm PBS (250 µl) to wash the migrated cells on the bottom of the membrane. Fifty µl of PBS was added to the top chamber before doing a second clean with a cotton swab. The PBS in the top chamber was aspirated by vacuum before the chamber was moved to a new receiver plate containing warmed 125 µl of a trypsin solution (Corning, cat # 25-053-C1). To detach the migratory cells from the bottom of the membrane, plates were incubated at 37 °C for 12 minutes, rocked for 5 minutes at room temperature, and tapped on each side. Trypsinized cells were rapidly transferred to a new white, opaque-bottom 96-well plate (Opti-Plate, Perkin Elmer, cat# 6005290) and mixed with an equal volume of CellTiterGlo (Promega, cat# G7570) diluted 1:3 in 10% FBS medium to estimate cell abundance. The mixture was shaken for 2 minutes and incubated for 10 minutes at room temperature before reading total luminescence in a Biotek Synergy H1 plate reader. Luminescent counts in the condition without FBS in the bottom chamber was subtracted from all conditions, and the resulting counts were normalized to counts in the DMSO-only control (100%) as a measure of migration. In compound dose dependence experiments, values of 3 technical replicates (wells) were averaged in each independent experiment.

**Cell viability assay**

Cells were seeded in 96-well culture plates as follows (number of cells per well in parenthesis): MDA-MB-231 (7,500); MCF-7 (12,500); MCF-10A (7,500); Hs-578T (7,500); BT-549 (7,500); HeLa (7,500). Each well contained 100 µl of the complete medium for each cell line described in "*Cell culture and establishment of cell lines*". After an overnight growth period, the medium was aspirated and replaced with 100 µl of the appropriate medium containing the final concentration of compounds indicated in the figures or figure legends or vehicle control (DMSO). DMSO concentration was equalized for all conditions in each experiment to 0.5% (v:v). After 24 hours of incubation with compound, the medium was aspirated and replaced with 100 µl of CellTiterGlo (Promega, cat# G7570) diluted with medium (1:5 ratio) that had been equilibrated to room temperature. The mixture was shaken for 2 minutes and incubated for 10 minutes at room temperature before reading total luminescence in a Biotek Synergy H1 plate reader. Luminescent counts detected in wells containing only medium were subtracted from all conditions, and the resulting counts were normalized to counts in the DMSO-only control (100%) as a measure of cell abundance. In compound dose dependence experiments, values of 3 technical replicates (wells) were averaged in each independent experiment.

**Nuclear Magnetic Resonance (NMR)**

All NMR data were measured on Bruker AVANCE 800 spectrometers equipped with cryogenically cooled triple resonance z-gradient probes. Proton chemical shifts were referenced to internal 2,2-dimethyl-2-silapentane-5-sulfonate (DSS, 0.00 ppm), and $^{13}C$ and $^{15}N$ chemical shifts were indirectly referenced to DSS (30). The NMR Spectra were processed with TopSpin (Bruker) and analyzed with Sparky (31). Gαi3 spectra were recorded at 30 °C on 400 µl samples in 5 mm Shigemi NMR tubes (without plunger) containing $^{2}H$-$^{13}C$-$^{15}N$ –Gαi3 in 10 mM HEPES pH 7.0 with 10 mM $MgCl_2$, 5 mm DTT, 0.01 % $NaN_3$ and 5% $^{2}H_2O$ with either 300 µM GDP or 300 µM GTPγS. The protein samples were prepared from a frozen stock solution of Gαi3 in 10 mM HEPES pH 7.0, 150 mM NaCl, 1 mM DTT and 20 µM GDP) by three cycles of 4-fold dilution (into 10 mM HEPES pH 7.0, 10 mM $MgCl_2$, 5 mm DTT, and 300 µM GDP or 300 µM GTPγS) and concentration by ultrafiltration using 10 kDa cut-off membranes. The NMR samples were prepared by addition of small volumes of concentrated stocks of $NaN_3$,$^{2}H_2O$ and DSS. The assignment of the NMR signals in the $^{1}H$-$^{15}N$ TROSY of Gαi3-GDP was done based on the Biological Magnetic Resonance Data Bank (BMRB) entry 19015 (32) corrected by adding 0.09 and -1.10 ppm to the $^{1}H$ and $^{15}N$ chemical shifts, respectively (2). The assignment of the NMR signals in the $^{1}H$-$^{15}N$ TROSY of Gαi3-GTPγS was done based on the BMRB entry 18103 (32) corrected by adding -0.05 and 0.58 ppm to the $^{1}H$ and $^{15}N$ chemical shifts, respectively (in this case, the BMRB chemical shifts were measured at pH 6.5, which may explain in part the difference). Titrations were done by the step wise addition of small volumes of a 10 mM solution of IGGi-11 in 50% aqueous DMSO (Gαi3-GDP) or a 100 mM solution of IGGi-11 in pure DMSO (Gαi3-GTPγS). The accumulated amount of DMSO added to the protein sample at the last titration point was less than 2 %, too low to significantly affect the chemical shift of the protein amide protons (33). The assignment of the NMR signals in the $^{1}H$-$^{15}N$ TROSY of Gαi3 bound to IGG1-11 was based on the nearest neighbor approach along the titrations. The Chemical Shift Perturbations were computed as the weighted average distance between the backbone amide $^{1}H$ and $^{15}N$ chemical shifts in the free and bound states, as described (34). For those residues with no assigned signal in the spectrum of free protein or without a reliable assignment in the bound protein, no chemical shift perturbation could be calculated and classified as "no data" in the corresponding figure. For some residues with weak signals in the spectrum of free Gαi3, the intensity decreased in the bound form below three times the level of the noise, beyond recognition as reliable signals, and are labeled as such in the corresponding figure.

**Dose-dependence fluorescence polarization (FP) assay**

Unless otherwise indicated in the figures or figure legends, this assay was carried out with rat His-Gαi3. The composition of the assay buffer was 10mM HEPES, 10mM $MgCl_2$, 0.0004% (v:v) NP-40, 5mM DTT, and 300 µM GDP, pH 7 with the exception of experiments shown in **Fig. S6**, which were done in buffer 50 mM Tris, 100 mM NaCl, 10 mM $MgCl_2$, 5 mM EDTA, 0.04% (v:v) NP-40, 30 µM GDP, 1 mM DTT, pH 7.4. In Figure

S6, the conditions with peptide KB-1753 also contained 30 µM $AlCl_3$ and 10 mM NaF (to generate the $AlF_4^-$ species that permits the association of Gαi3 with this peptide (3)). Five µl of Gα protein were added to black 384-well plates (OptiPlate-384F, Perkin Elmer cat#6007270), mixed with 10 µl of compound, and incubated for 10 minutes at room temperature. Then, 5 µl of FITC-labeled peptide (GIV, R12 GL, KB-1753) were added to each well, and incubated again for 10 min at room temperature protected from light before reading fluorescence. Fluorescence polarization (Ex 485 ± 10 nm/Em 528 ± 10 nm) was measured every 2 min for 30 min at room temperature in a Biotek H1 synergy plate reader to ensure that the signals were stable in time. Fluorescence polarization at different times was averaged for all subsequent calculations. The final concentration of Gα protein was 1 µM and the final concentration of peptides was 25 nM. Compounds were at the concentrations indicated in the figures or figure legends, and the concentration of DMSO was equalize to 1% (v:v) across all conditions, including negative controls without compound. Conditions containing FITC-labeled peptides but no G-protein were included to determine the basal FP levels to be subtracted from all conditions, which were subsequently normalized using the DMSO-only controls as 100% reference. Results were fitted to a one-site competitive binding model inhibition curve to calculate the $IC_{50}$ values using Prism (GraphPad). $IC_{50}$ values were converted to inhibitor constants ($K_i$) (35) by using previously determined equilibrium dissociation constants ($K_D$) (2, 3).

**Computational docking of IGGi-11 on Gαi3**

IGGi-11 was docked on a previously described (2) model of GIV-bound Gαi3 (coordinates deposited at www.modelarchive.org; 10.5452/ma-ayq5v). After removing GIV from the model, the structure of IGGi-11 was docked using ICM version 3.8–3 (Molsoft LLC., San Diego, CA) by focusing on a region in the vicinity of Gαi3 residues K35, K197, G217, S252, W258, and R313. IGGi-11 was treated as a fully flexible ligand in a rigid receptor simulation within continuous dielectric solvent using internal coordinate mechanics (36, 37). Force fields and energy potentials are determined with the modified Merck Molecular Force Field 94 (MMFF94) (38) and the Empirical Conformation Energy Program for Peptides (ECEPP/3) (39) for small molecules and proteins, respectively. Internal force field energy, receptor-ligand hydrogen bonding, hydrophobic energy, conformational entropy loss, and solvation energy change were used for scoring and ranking of docking poses. The maximum van der Waals repulsion was set to 4.0. The receptor maps had a grid size of 0.5 angstroms. No explicit waters were included in the simulation. Structure images were rendered with ICM (Molsoft) or PyMol (Schrodinger).

**GST pull-down assay**

Assessment of protein-protein binding using GST pull-down assays was carried out as described previously (4, 22, 40) with minor modifications. Briefly, 3 µg of GST-GIV (aa 1671-1755), GST-RGS4, or GST were immobilized on glutathione agarose beads (ThermoFisher#16100) for 90 min at room temperature in PBS. Beads were washed twice with PBS and resuspended in 250 µl of binding buffer (50 mM Tris-HCl, pH 7.4, 100 mM NaCl, 0.04% (v:v) NP-40, 10 mM $MgCl_2$, 5 mM EDTA, 1 mM DTT, 30 µM GDP), and supplemented with test compounds (100 µM) or an equivalent volume of DMSO. After addition of 50 ng of (~ 5 nM) of rat His-Gαi3 purified, tubes were incubated for 4 hr at 4 °C with constant rotation. After this incubation, beads were washed four times with 1 ml of wash buffer (4.3 mM $Na_2HPO_4$, 1.4 mM $KH_2PO_4$, pH 7.4, 137 mM NaCl, 2.7 mM KCl, 0.1% (v/v) Tween-20, 10 mM $MgCl_2$, 5 mM EDTA, 1 mM DTT, 30 µM GDP). For the experiments testing binding of G proteins to RGS4, all buffers were supplemented with 30 µM $AlCl_3$ and 10 mM NaF to load Gαi with GDP·$AlF4^-$. Resin-bound proteins were eluted by boiling for 5 min in Laemmli sample buffer, and proteins were separated by SDS-PAGE and immunoblotted with antibodies as indicated under "*Cell signaling stimulation, cell lysis, and immunoblotting*".

**Isothermal titration calorimetry (ITC)**

These experiments were carried out at 25 °C using a MicroCal iT200 system (Malvern Panalytical, Malvern, UK). IGGi-11 was diluted from a 100 mM stock in DMSO to a final concentration of 1 mM in assay buffer (10 mM HEPES, 10 mM $MgCl_2$, 1 mM TCEP, 300 µM GDP, pH 7.0). Experiments were performed by injecting 2 µl of this compound solution into a 200 µl solution containing 50 µM human His-Gαi3 (WT or mutants) supplemented with 1% DMSO in the sample cell. A total of 19 sequential injections were performed with a spacing of 150 s and a reference power of 9 µcal/s. For each measurement session, a control experiment to estimate heat of dilution was carried by compound titration into buffer without protein. The heat of dilution was subtracted from each compound-protein titration, and the binding isotherms were plotted and analyzed using Origin Software (MicroCal Inc., USA). Data were fit to a one-site binding model (N = 1). Each protein was analyzed at least twice with similar results.

**Bioluminescence Resonance Energy Transfer (BRET) measurements in isolated membranes**

HEK293T cells were seeded on 6-well plates (~400,000 cells/well) coated with 0.1% gelatin and transfected 24 hr later using the calcium phosphate method. For experiments using the free Gβγ biosensor system, all conditions received the following amounts of plasmid per dish: 1.2 µg for VC-$G\beta_1$, 1.2 µg for VN-$G\gamma_2$, and 0.3 µg mas-GRRK3ct-Nluc. In addition, plasmids for the following combinations of Gα subunit and GPCR were co-transfected: 6 µg for Gαi3 and 1.2 µg for $\alpha 2_A$-AR; 6 µg for Gαo and 1.2 µg for $\alpha 2_A$-AR; 6 µg for Gαq-HA and 1.2 µg for M3R; and 6 µg for Gα13 and 1.2 µg for PAR1. For experiments aimed at detecting Gαi-GTP, cells were transfected with the following amounts of plasmid DNA per dish: 1.2 µg for $\alpha 2_A$-AR, 0.3 µg for mas-KB1753-Nluc, 3 µg for Gαi3-YFP (b/c loop). Approximately 18-24 hr after transfection, cells were scraped in PBS and pelleted at 550 x g for 5 min. Pellets from one 10-cm dish were resuspended in 250 µl of ice-cold homogenization buffer (10 mM HEPES, 250 mM sucrose, 10 mM KCl, 1.5 mM $MgCl_2$, 1 mM DTT, pH 7.4) supplemented with a protease inhibitor cocktail (Sigma, cat#8820). All subsequent steps were carried out in ice or at 4 °C. Cells were homogenized by 30 passages though a 30-gauge needle and subsequently centrifuged at 1,000xg for 10 min. The pellet was discarded and the supernatant transferred to a new tube (Thermo Scientific, cat#314352) and centrifuged in a TLA-55 fixed angle rotor in a Beckman Coulter Optima MAX-E tabletop centrifuge for 45 min at 100,000xg. The supernatant was aspirated, and the pellet was resuspended in 250 µl of homogenization buffer by pipetting and syringing. This fraction containing isolated cell membranes was stored as single-use 10 µl aliquots at -80 °C. Both endpoint and kinetic BRET measurements were carried out in a final volume of 100 µl in 96-well plates. Briefly, 5 µl of isolated cell membrane fraction was mixed with 85 µl of assay buffer (140 mM NaCl, 5 mM KCl, 1 mM $MgCl_2$, 1 mM $CaCl_2$, 0.37 mM $NaH_2PO_4$, 24 mM $NaHCO_3$, 10 mM HEPES, 0.1% (w:v) glucose, 1mM DTT, 0.002% (w:v) BSA, pH 7.4, supplemented with protease inhibitor cocktail at one-tenth of the recommended concentration for cell lysates) containing the required amounts of IGGi-11 to achieve the final concentration of compound indicated in the figure or figure legends. The concentration of DMSO was equalized to 1% (v:v) across all conditions. In the experiments to assess the association of Gβγ with Gα in the absence of GPCR stimulation, the buffer was supplemented with 300 µM GDP (Alfa Aesar #J61646), except for the control condition containing GTPγS, in which GDP was replaced by 300 µM GTPγS (Sigma G8634). As an additional control condition in these experiments, the buffer was supplemented with 25 µM of the R12 GL peptide, a Gαi binding peptide previously reported to disrupt its association with Gβγ (41). In the experiments to assess the dissociation of Gβγ form Gα or for the formation of Gαi3-GTP upon GPCR stimulation, the buffer was supplemented with 300 µM GTP (MB Bioscience #151216), except for the control conditions containing GTPγS, in which GTP was replaced by 300 µM GTPγS (Sigma G8634). GPCR agonists were used as follows: 1 µM brimonidine (Ark Pharm, cat# AK-3579) for experiments with Gi, 0.1 µM brimonidine for experiments with Go, 100 µM carbachol (Acros Organics cat# AC-10824) for experiments with Gq, and 30 µM Thrombin Receptor Activator Peptide 6 (TRAP-6, Anaspec cat# AS-24190) for experiments with G13. For all endpoint experiments, membrane/compound mixtures were incubated for 3 min at room temperature before adding 10 µl of coelenterazine 400a to obtain a final concentration of 5 µM. Two minutes after the addition of coelenterazine 400a, luminescence signals were measured in a POLARstar OMEGA plate reader (BMG Labtech) at 28 °C. Luminescence was measured at 450 ± 40 and 535 ± 15 nm, and BRET was calculated as the ratio between the emission intensity at 535 ± 15 nm divided by the emission intensity at 450 ± 40 nm. Ratios determined from three consecutive measurements spaced by 36 s were averaged. Results were presented as the BRET change relative to an untreated condition (no IGGi-11 and not GPCR agonist). The procedures were similar for kinetic BRET measurements, except that luminescence signals were measured every 0.24 s for the duration of the experiment, and that brimonidine (0.1 µM) and yohimbine (100 µM) were injected into the wells during the measurements as indicated in the figures. Results for the kinetic BRET measurements were presented as the BRET change relative to the baseline signal (the average BRET ratio in the 30 s before agonist stimulation).

**Adenylyl cyclase activity in isolated membranes**

Two million HEK293T cells were seeded on gelatin coated 10 cm dishes. Eighteen hours later, cells were transfected with 6 μg of a plasmid DNA encoding human AC5-YFP using the calcium phosphate method. Twenty-four hours after transfection, cells were scraped in PBS and pelleted by centrifugation at 550×g for 5 minutes. Cell pellets were re-suspended in homogenization buffer and membranes isolated as described in "*Bioluminescence Resonance Energy Transfer (BRET) measurements in isolated membranes*". Membrane pellets were resuspended in 250 µl of homogenization buffer, and protein content quantified by Bradford. Aliquots were stored at -80 °C until their use in experiments. Adenylyl cyclase activity in membranes was determined by the production of cAMP under different conditions. Reactants were mixed on ice in a final volume of 40 µl of assay buffer (50 mM HEPES, 2 mM $MgCl_2$, 1 mM EDTA, 0.5 mg/mL BSA, pH 8.0) as follows. All conditions were done in duplicate. Four µl of IGGi-11 or vehicle (DMSO) were mixed with 8 µl of myr-Gαi1-GTPγS (or the same volume of buffer for conditions without Gαi1) and incubated for 10 minutes before the addition of 2 µg of AC5-expressing membrane protein in a volume of 8 µl. Next, 8 µl of Gαs-GTPγS, forskolin or buffer were added and tubes were incubated on ice for 10 minutes. Reactions were started by adding pre-warmed ATP and $MgCl_2$ solution in a volume of 4 µl and rapidly transferring the tubes to a heat block at 30 °C for 10 minutes. Reactions were stopped by heating tubes at 95°C for 5 minutes, then centrifuged and an aliquot from the supernatant was taken to quantify cAMP using the LANCE cAMP kit (Perkin Elmer, cat#AD0262) according to the manufacturer protocol. The final concentrations of reactants were: IGGi-11, 100 µM; DMSO, 0.1 % (v:v); myr-Gαi1-GTPγS, 1 µM, Gαs-GTPγS, 0.1 µM; forskolin (Tocris, cat#1099), 10 µM; ATP, 1 mM; $MgCl_2$, 5 mM. myr-Gαi1 and Gαs were loaded with GTPγS by incubating them at 30 °C with 150 µM GTPγS in buffer (20 mM Tris-HCl, 20 mM NaCl, 1 mM $MgCl_2$, 1 mM DTT, 5 % glycerol (v:v) for 3 hours or 45 minutes, respectively. Time-resolved fluorescence measurements to quantify cAMP were done on a TECAN Infinite M1000 plate reader in white 384-well ProxiPlates (Perkin Elmer, cat#6008280). Specific activity was expressed as pmol cAMP / min / mg membrane after background subtraction of the cAMP signal obtained in the absence of membranes. Values of 2 duplicates were averaged in each independent experiment.

**Steady-state GTPase assay**

This assay was performed as described previously (4, 6, 22, 40) with minor modifications. Briefly, purified human His-Gαi3 WT (500 nM) or human His-Gαi1 R178M/A326S (Gαi1$^{RM/AS}$, 50 nM) was pre-incubated for 15 min at 30 °C in assay buffer (20 mM Na-HEPES, 100 mM NaCl, 1 mM EDTA, 2 mM $MgCl_2$, 1 mM DTT, and 0.05% (w:v) $C_{12}E_{10}$, pH 8) with the concentrations of IGGi-11, His-GIV-CT, R12 GL peptide, or His-RGS4 indicated in the figures or figure legends. GTPase reactions were initiated at 30 °C by adding an equal volume of assay buffer containing 1 µM [γ-$^{32}$P]GTP (~50 c.p.m./fmol). Duplicate aliquots (25 µL) were removed at 10 min and reactions stopped with 975 µl of ice-cold 5% (w:v) activated charcoal in 20 mM $H_3PO_4$, pH 3. Samples were then centrifuged for 10 min at 10,000xg, and 500 µl of the resultant supernatant were scintillation-counted to quantify the amount of [$^{32}$P]Pi released. Background [$^{32}$P]Pi detected at 10 min in the absence of G protein was subtracted from each reaction. Background counts were <5% of the counts detected in the presence of G proteins. Results were calculated as relative to the activity of the G-protein alone (% of control).

**GTPγS-BODIPY binding assay**

GTPγS-BODIPY (Life Technologies, cat#G22183) was diluted in 100 µl of assay buffer (20 mM Na-HEPES, 100 mM NaCl, 1 mM EDTA, 2 mM $MgCl_2$, 1 mM DTT, 0.05% (w:v) $C_{12}E_{10}$, pH 8) to a final concentration of 50 nM in the presence of 30 µM IGGi-11, 30 µM NF023 (a positive control for inhibition of GTP binding to Gαi proteins (42)), or an equivalent amount of DMSO (0.1% v:v). After approximately 15 min incubation at 28 °C, fluorescence measurements were carried out at the same temperature by exciting at 485 nm and detecting emission at 535 ± 30 nm in a POLARstar OMEGA plate reader (BMG Labtech) every 30 seconds. Purified human His-Gαi3 (200 nM) was added in real time during the measurements as indicated in the figures.

**Membrane permeability and hydrolytic processing of compounds**

Membrane permeability of IGGi-11 or IGGi-11me was assessed using a parallel artificial membrane permeability assay (PAMPA) kit (BioAssay Systems, cat#PAMPA-096) following the manufacturer's instructions with minor modifications. Briefly, an artificial membrane was reconstituted over a porous support that separated the donor (upper) from an acceptor (bottom) well with a 4% (w:v) lecithin solution in dodecane. 100 µl of a solution containing compound diluted in PBS buffer (pH 7.4) at a final concentration of 20 µM in 5% (v:v) DMSO were added to the upper well, and 220 µl of 5% (v:v) DMSO in PBS buffer were added to the bottom well. After incubation for 6 hours at 37 °C, 90 µl or 200 µl were taken from the upper or bottom well, respectively, and transferred to microcentrifuge tubes (Olympus 1.7ml Microtube Cat# 24-282LR) that were stored at -80 °C. All conditions were done in duplicate. Samples were dried in a speed vacuum centrifuge and reconstituted with 0.1% (v:v) formic acid in water (90 µl or 30 µl for sample form the upper or the bottom wells, respectively) by vortexing. Samples were analyzed by LC-MS/MS. Chromatography was performed using a Waters Acquity CSH™ Phenyl-Hexyl 1.7µM 2.1 x 50mm column and acetonitrile/water as the mobile phase, and a Sciex API 4000 triple quadrupole mass spectrometer with an ESI source was used in positive mode with a full MS scan at 55.0-1000 m/z. Compound abundance was estimated from the area under the spectral curve (AUC) in FreeStyle 1.8 SP1 software (ThermoFisher). For the experiments assessing the hydrolytic processing of IGGi-11me into IGGi-11, samples were analyzed the same way by LC-MS/MS with some modifications in the procedure for sample preparation. Briefly, 90 µl of 20 µM IGGi-11me diluted in PBS with 5% (v:v) DMSO was incubated in the presence or absence of 25 µg of a cytosolic fraction of MDA-

MB-231 cells (43) for 2 hours at 37 °C or not incubated at all (time 0). All conditions were done in duplicate. Samples were extracted by mixing with 8 volume parts of an organic solution (8:1:1 Acetonitrile:Methanol:Acetone), incubation at 4 °C for 30 minutes, and centrifugation at 15,000xg for 15 min at 4 °C. The extracted clear supernatant was transferred to a clean tube and dried in a speed vacuum centrifuge. Samples were reconstituted with 90 µl of 0.1% (v:v) formic acid in water by vortexing before injection into the LC-MS/MS instrument.

**Cell signaling stimulation, cell lysis, and immunoblotting**

Two-hundred thousand MDA-MB-231 cells or 250,000 HeLa cells per well were seeded on 6-well plates. Twenty-four hours after seeding, cells were incubated overnight (~16 h) before EGF (Recombinant Human Epidermal Growth Factor, GoldBiotechnnology, cat#1150-04-100) stimulation with the concentrations of IGGi-11me indicated in the figures or figure legends or a matching amount of DMSO (0.25% v:v) in the presence of 0.5% (v:v) or 0.2% (v:v) FBS for MDA-MB-231 or HeLa cells, respectively. For experiments using SDF-1α (R&D Systems, 350-NS/CF) instead of EGF for stimulation, cells were incubated overnight before stimulation in the absence of FBS. In some cases, cells were also pre-incubated overnight with pertussis toxin (PTX, 100 ng/ml, List Biological Laboratories #179A). Cells were stimulated with EGF or SDF-1α as indicated in the figures or figure legends by adding concentrated stocks to the wells. Stimulation reactions were stopped by rapidly washing with ice-cold PBS three times and adding 100 µl of lysis buffer (20 mM HEPES, 5 mM $Mg(CH_3COO)_2$, 125 mM $K(CH_3COO)$, 0.4% (v:v) Triton X-100, 1 mM DTT, 10 mM β-glycerophosphate, and 0.5 mM $Na_3VO_4$, pH 7.4) supplemented with a protease inhibitor cocktail (Sigma, cat# S8830) per well before harvesting by scraping. Whole cell lysates were cleared by centrifugation (10 min at 14,000 × g, 4°C, and then quantified by Bradford (Bio-Rad, cat#5000205), and boiled in Laemmli sample buffer for 5 min before protein separation by SDS-PAGE and electrophoretic transfer to PVDF membranes (EMD Millipore, cat#IPFL00010) for 2 hr. PVDF membranes were blocked with TBS supplemented with 5% non-fat dry milk for 1 hr, and then incubated sequentially with primary and secondary antibodies. Primary antibody species, vendors, and dilutions were as follows: GIV (Rabbit, Santa Cruz Biotechnology, sc-133371); total Akt (Mouse, Cell Signaling Technologies, 2920), 1:2000; phosphorylated Akt (S473) (Rabbit, Cell Signaling Technologies, 9271), 1:1000; Gαi3 (Rabbit, Aviva, #OAAB19207), 1:250; Gαi1/2 (Rabbit, Sigma, 06-236) 1:250; β-actin (mouse, LiCor Biosciences, 926-42212), 1:2500; GST (Rabbit, Sigma, G7781), 1:2500; His (mouse, Sigma, H1029), 1:2500. Secondary antibodies (goat anti-rabbit conjugated to AlexaFluor 680 (Life Technologies, #A-21077) or goat anti-mouse conjugated to IRDye 800 (LI-COR Biosciences, #926–32210) were used at 1:10,000. Infrared imaging of immunoblots was performed according to manufacturer's recommendations using an Odyssey CLx infrared imaging system (LI-COR Biosciences). Akt activation was determined by calculating the phospho-Akt (pAkt)/ total-Akt ratio and normalizing it to the maximum activation in each experiment (percentage of maximal response). Images were processed using the ImageJ software

(NIH) and assembled for presentation using Photoshop and Illustrator software (Adobe). The same protocol of cell lysis and immunoblotting was followed to detect the expression of proteins in any other cell line under steady state culture condition or for the detection of proteins in GST pulldown experiments.

**Bioluminescence Resonance Energy Transfer (BRET) measurements in HeLa cells**

BRET experiments with already generated HeLa cells stably expressing biosensor constructs for Gαi-GTP (Gαi*-BERKY3) or for free Gβγ (Gβγ-BERKY3) were carried out as described previously (29).

**Matrigel cultures**

MDA-MB-231 or MCF10A cells were cultured on top of Matrigel as described previously (28, 44) with minor modifications. Briefly, 25 µl of ice-cod Matrigel (growth factor reduced, Corning, #356231) were spread as a thick layer on the bottom of 96-well plates (Thermo 167008) and allowed to solidify at 37 °C. Two-thousand and five hundred cells were seeded on the Matrigel-coated wells in a volume of 100 µl of their regular growth medium supplemented with 2% (v:v) Matrigel. Six hours after cell seeding, the medium was replaced with 100 µl of the same medium supplemented with 100 µM IGGi-11me or vehicle (0.5% DMSO). All conditions were done in duplicates. Medium was replaced every other day by fresh medium without compound. Cell abundance was estimated on days 0, 2, 4, 5, and 7 using CellTiter-Glo® (Promega, G7570) as described in "*Cell viability assays*" section. Relative Luminescence Unit (RLU) values of the technical replicates (wells) were averaged in each independent experiment. For imaging of Matrigel cultures, cells were grown under equivalent conditions but in 8-well glass bottom chambers (Cellvis, cat# C8-1.5H-N). Briefly, 5,000 cells were seeded on wells coated with 40 µl of Matrigel in a volume of 200 µl. Images were acquired 7 days after seeding by phase-contrast microscopy with a Zeiss Axio Observer Z1microscope equipped with a camera (C10600/Orca-R2; Hamamatsu Photonics) using 10x (NA= 0.45; working distance 2 mm) or 20x (NA= 0.8; working distance 0.55 mm) objectives. Images were acquired and processed using Zeiss' ZEN software and assembled for presentation using Photoshop and Illustrator software (Adobe).

**Mouse xenograft and tail vein injection**

Half a million MDA-MB-231 cells stably expressing fluc2-myc were seeded on 10 cm dishes. Twenty-four hours after seeding, medium was replaced with DMEM supplemented with 0.2% FBS and 100 µM IGGi-11me or vehicle (0.5% DMSO) and dishes placed in the cell incubator overnight. After this, cells were detached by incubation with warm citric saline solution (135 mM KCl, 15 mM sodium citrate, pH 7.2) or Versene solution (Gibco, #15040066), washed three times with serum-free DMEM, and resuspended in serum-free DMEM at a concentration of $2 \times 10^7$ cells/ml for flank injections or $1 \times 10^6$ cells/ml for tail vein injections. $2 \times 10^6$ cells were injected subcutaneously in the hind flank of mice, or $2.5 \times 10^5$ cells in the tail vein. For all studies, ~8 week-old female NCr nu/nu athymic nude mice (Taconic Bioscience, cat#NCRNU-F) were used. Tumor cells were visualized using whole-body bioluminescence imaging (BLI) using an IVIS® Spectrum In Vivo Imaging System (Perkin Elmer) 3-5 minutes after intraperitoneal injection of 150 mg/kg D-luciferin potassium salt (GoldBiotechnology, cat#LUCK-1G) dissolved in PBS (Corning, 21-040-CV) and filtered through a 0.45-μm surfactant-free cellulose acetate (SFCA) membrane filter. Photon count per second per square centimeter per steradian (p/sec/cm$^2$/sr) values and images were acquired using Living Images software (Perkin Elmer). At the end of experiments, flank injected tumors were removed and photographed. All individual images were processed using Living Images software and assembled for presentation using Photoshop and Illustrator software (Adobe). All animal procedures were approved by the IACUC of Boston University under protocol PROTO201800258.

FIGURE S1

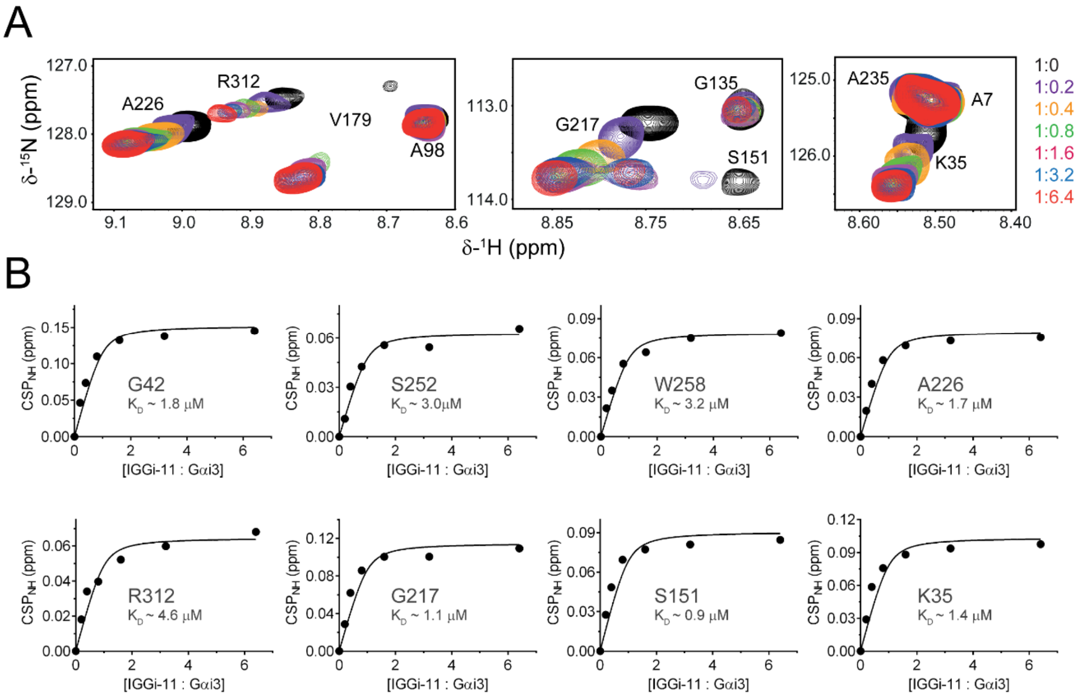


**Figure S1. Gαi3 NMR signal perturbations upon IGGi-11 titration. (A)** Overlay of $^{1}H$-$^{15}N$- TROSY spectra of $^{2}H$,$^{13}C$,$^{15}N$-Gαi3-GDP after addition of increasing amounts of IGGi-11. Colors correspond to the molar Gαi3:IGGi-11 ratios indicated on the right. **(B)** Plots of the measured chemical shift perturbation (CSP) values of selected Gαi3 residues fitted to a single-site binding model to estimate equilibrium dissociation constants.

FIGURE S2

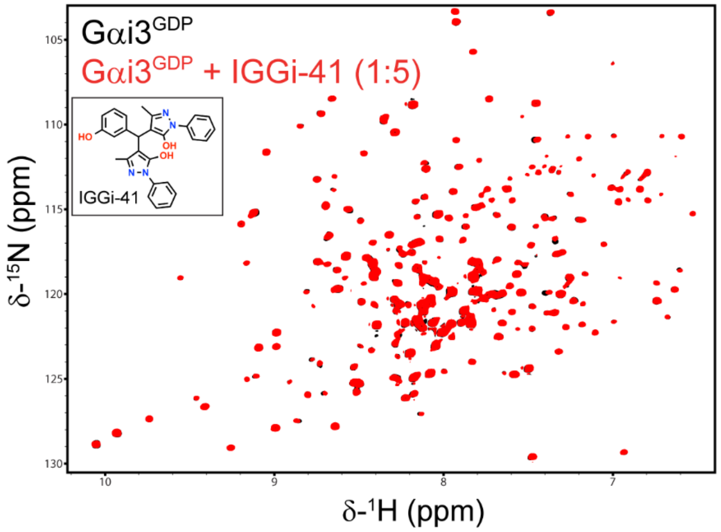


**Figure S2. IGGi-41 does not induce NMR signal perturbations on Gαi3. (A)** Overlay of $^{1}H$-$^{15}N$ TROSY spectra of $^{2}H$,$^{13}C$,$^{15}N$-Gαi3-GDP in the absence or presence of IGGi-41 in five-fold molar excess, showing minimal perturbations caused by the compound.

FIGURE S3

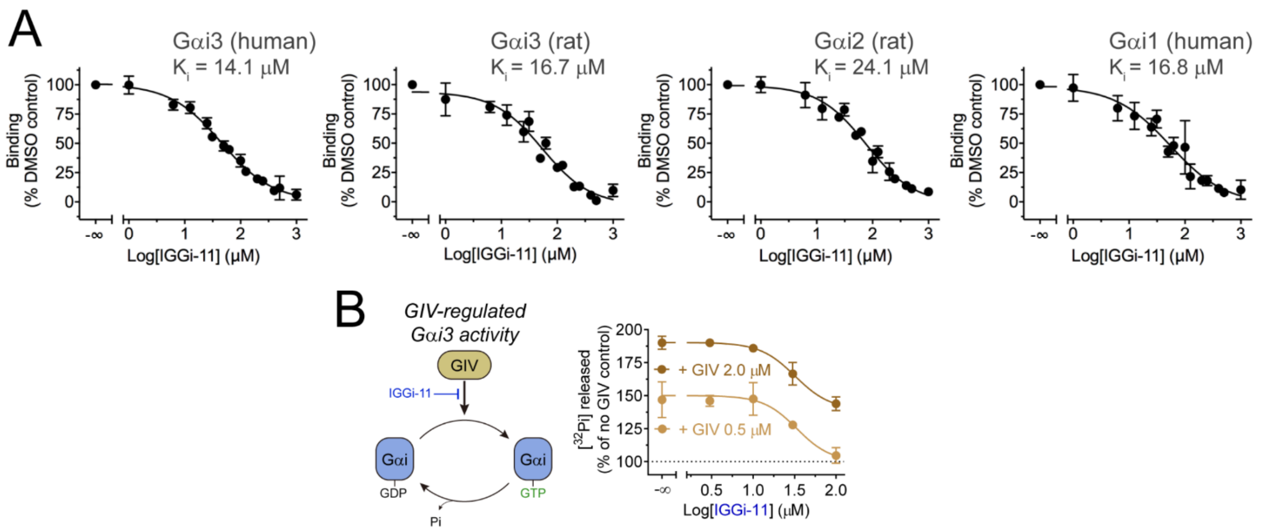


**Figure S3. IGGi-11 inhibits binding of GIV to different Gαi subunits, and blocks GIV-mediated G protein activity regulation *in vitro*. (A)** Quantification of GIV binding to different Gαi proteins by fluorescence polarization in the presence of increasing concentrations of IGGi-11. $K_i$ values were determined from the curve fits as indicated in *Methods*. Results are expressed as mean ± SEM (*N* = 3). **(B)** IGGi-11 inhibits GIV-mediated stimulation of Gαi3 steady-state GTPase activity. The steady-state GTPase activity of Gαi3, which depends on the rate of nucleotide exchange, was determined in the presence of 0.5 or 2 μM GIV-CT and increasing concentrations (1–100 μM) of IGGi-11 by measuring the production of $[^{32}P]P_i$ from GTP[γ-$^{32}$P]. Results are expressed as % of the activity of Gαi3 in the absence of GIV. Mean ± S.E.M (*N* = 3).

### FIGURE S4

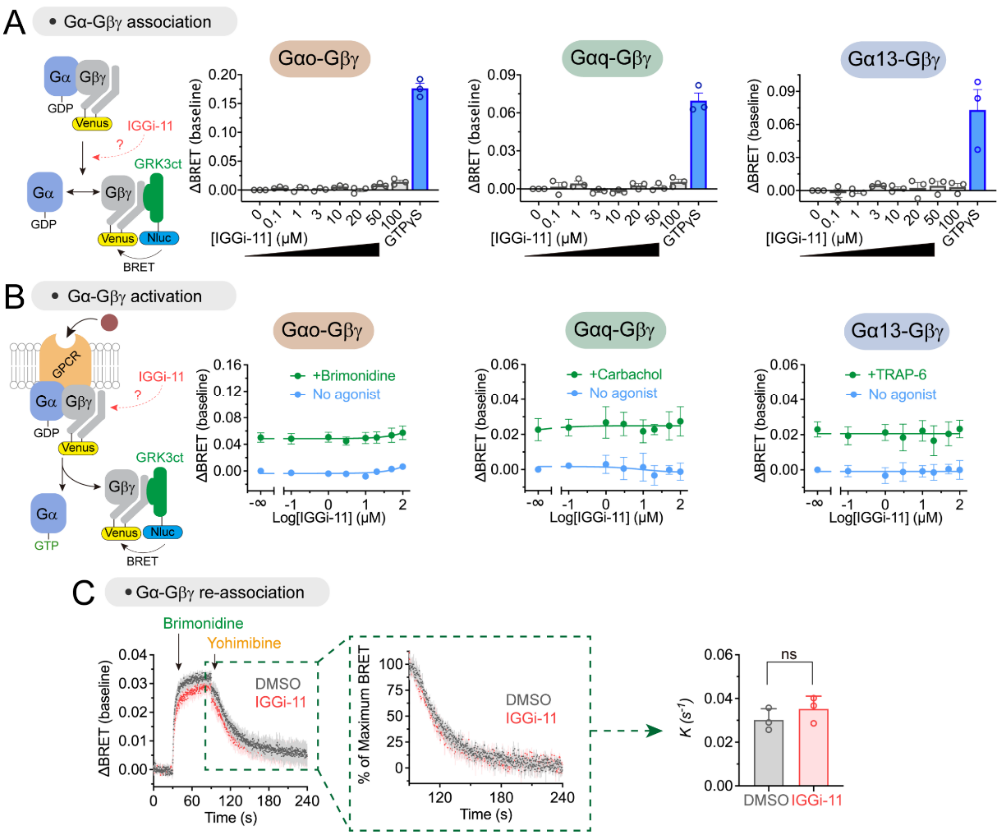


**Figure S4. Lack of effect of IGGi-11 on G-protein heterotrimer association, re-association, or coupling to GPCRs. (A)** IGGi-11 does not dissociate Gβγ from Gαo, Gαq or Gα13 in membranes isolated from HEK293T cells expressing a BRET-based biosensor for free Gβγ, whereas GTPγS (300 μM) does. **(B)** IGGi-11 does not affect GPCR-mediated activation of Go, Gq or G13 in membranes isolated from HEK293T cells as determined by the dissociation of Gα-Gβγ heterotrimers using BRET-based biosensors. The $\alpha 2_A$ adrenergic receptor, the M3 muscarinic receptor, or the PAR1 receptor were co-expressed for experiments with Go, Gq or G13, respectively. Membranes were treated with the indicated concentrations of IGGi-11 with (green) or without (blue) stimulation with a receptor agonist (1 μM brimonidine for Go, 100 μM carbachol for Gq, and 30 μM Thrombin Receptor Activator Peptide 6 (TRAP-6) for G13) for 2 minutes before BRET measurements. **(C)** IGGi-11 does not interfere with Gβγ re-association with Gαi3 upon termination of GPCR stimulation. Membranes isolated from HEK293T cells co-expressing Gαi3, the $\alpha 2_A$ adrenergic receptor, and the components of a BRET-based biosensor for free Gβγ were treated with brimonidine (agonist, 0.1 μM) and yohimbine (antagonist, 100 μM) as indicated during continuous kinetic luminescence measurements. Deactivation rates (*k*) were determined by fitting normalized deactivation data after antagonist addition to an exponential decay curve. All data are mean ± SEM (*N* = 3-4).

### FIGURE S5

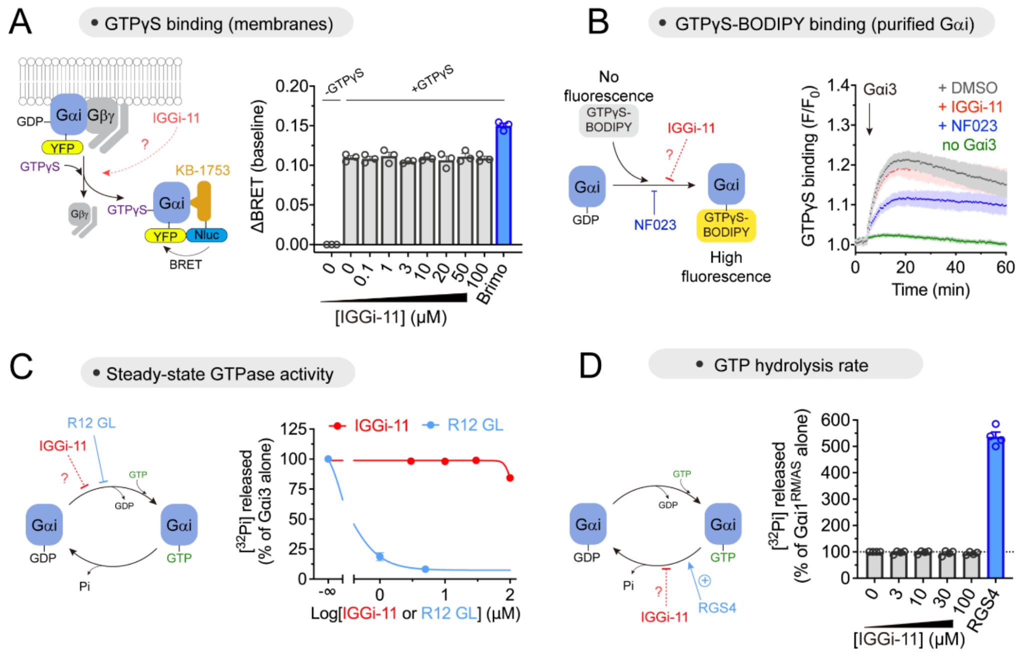


**Figure S5. Lack of effect of IGGi-11 on nucleotide handling by G-proteins. (A)** IGGi-11 does not affect spontaneous GTPγS binding to Gi3 as determined by BRET-based detection of Gαi3-GTP in isolated cell membranes. Membranes isolated from HEK293T cells expressing a BRET-based biosensor for Gαi3-GTP were incubated with GTPγS (300 μM) and the indicated concentrations of IGGi-11. GPCR stimulation with brimonidine (1 μM) increases GTPγS binding. Mean ± SEM (*N* = 3). **(B)** IGGi-11 does not affect spontaneous GTPγS binding to purified Gαi3 as determined by a fluorescent analog assay. Purified Gαi3 was added to a solution containing GTPγS-BODIPY and the increase in fluorescence caused by binding of the nucleotide to the G-protein monitored continuously in the presence of 30 μM IGGi-11 or NF023 (a positive control for inhibition of nucleotide binding by Gαi (42)), or DMSO (1 %) as the negative control. Mean ± SEM of 3 independent experiments. **(C)** IGGi-11 does not affect Gαi3 steady-state GTPase activity. The steady-state GTPase activity of Gαi3, which depends on the rate of nucleotide exchange, was determined in the presence of increasing concentrations (3-100 μM) of IGGi-11 or the GoLoco peptide R12 GL (1-5 μM, positive inhibition control), by measuring the production of [$^{32}$P]P$_i$ from GTP[γ-$^{32}$P]. Results are expressed as % of the activity of Gαi3 alone. Mean ± S.E.M (*N* = 3). **(D)** IGGi-11 does not affect GTP hydrolysis by Gαi. The steady-state GTPase activity of the Gαi1$^{RM/AS}$ mutant, which depends on the rate of nucleotide hydrolysis but not of nucleotide exchange (45), was determined in the presence of increasing concentrations (3-100 μM) of IGGi-11 or 0.8 μM of the GAP RGS4 (positive control for enhancement of nucleotide hydrolysis), by measuring the production of [$^{32}$P]P$_i$ from GTP[γ-$^{32}$P]. Results are expressed as % of the activity of Gαi1$^{RM/AS}$ alone. Mean ± S.E.M (*N* = 4).

FIGURE S6

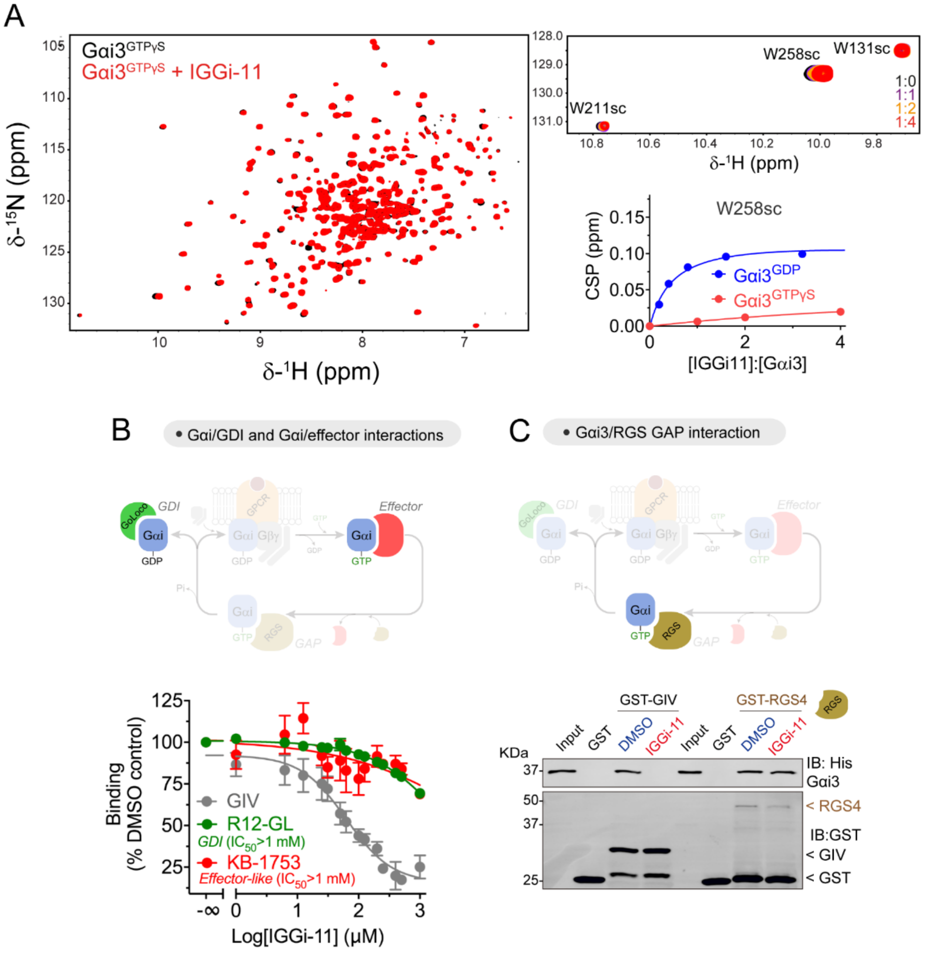


**Figure S6. IGGi-11 does not bind to GTP-bound Gαi3 or prevent Gαi3 binding to GDIs, GAPs or effectors. (A)** Overlay of $^{1}$H-$^{15}$N TROSY spectra of $^{2}$H,$^{13}$C,$^{15}$N-Gαi3-GTPγS in the absence or presence of IGGi-11 display minimal perturbations by the compound. A selected region containing signals for the side chains of W211 and W258, which undergo large perturbation in GDP-bound Gai3 in the presence of IGGi-11 (**Fig. 2**), is shown enlarged on the upper right. The lower right plot compares the chemical shift perturbation (CSP) values of the side chain of W258 in GDP- or GTPγS-bound Gαi3 upon IGGi-11 titration. **(B)** IGGi-11 inhibits GIV binding but does not inhibit the binding of a GoLoco GDI peptide or an effector-like peptide to rat Gαi3, while it blocks GIV binding. Binding of a GIV peptide, a GDI peptide corresponding to the GoLoco motif of RGS12 (R12-GL), or the effector-like peptide KB-1753 to Gαi3 was quantified by fluorescence polarization. GIV and R12-GL experiments were done in the presence of GDP, whereas KB-1753 experiments were done in the presence of GDP+$AlF_4^-$. Mean ± SEM ($N$ = 3). **(C)** IGGi-11 disrupts GIV-Gαi3 binding but not GAP-Gαi3 binding in pulldown assays. Gαi3 was incubated with glutathione agarose-bound GST-GIV (aa 1671-1755) or GST-RGS4 in the presence of IGGi-11 (100 μM) or DMSO (1%). After incubation and washes, bead-bound proteins were separated by SDS-PAGE and immunoblotted (IB) as indicated. Conditions with GIV contained GDP, whereas those with RGS4 contained GDP + $AlF_4^-$ to induce the formation of the transition state recognized by GAPs. One experiment representative of 3 independent repeats is presented.

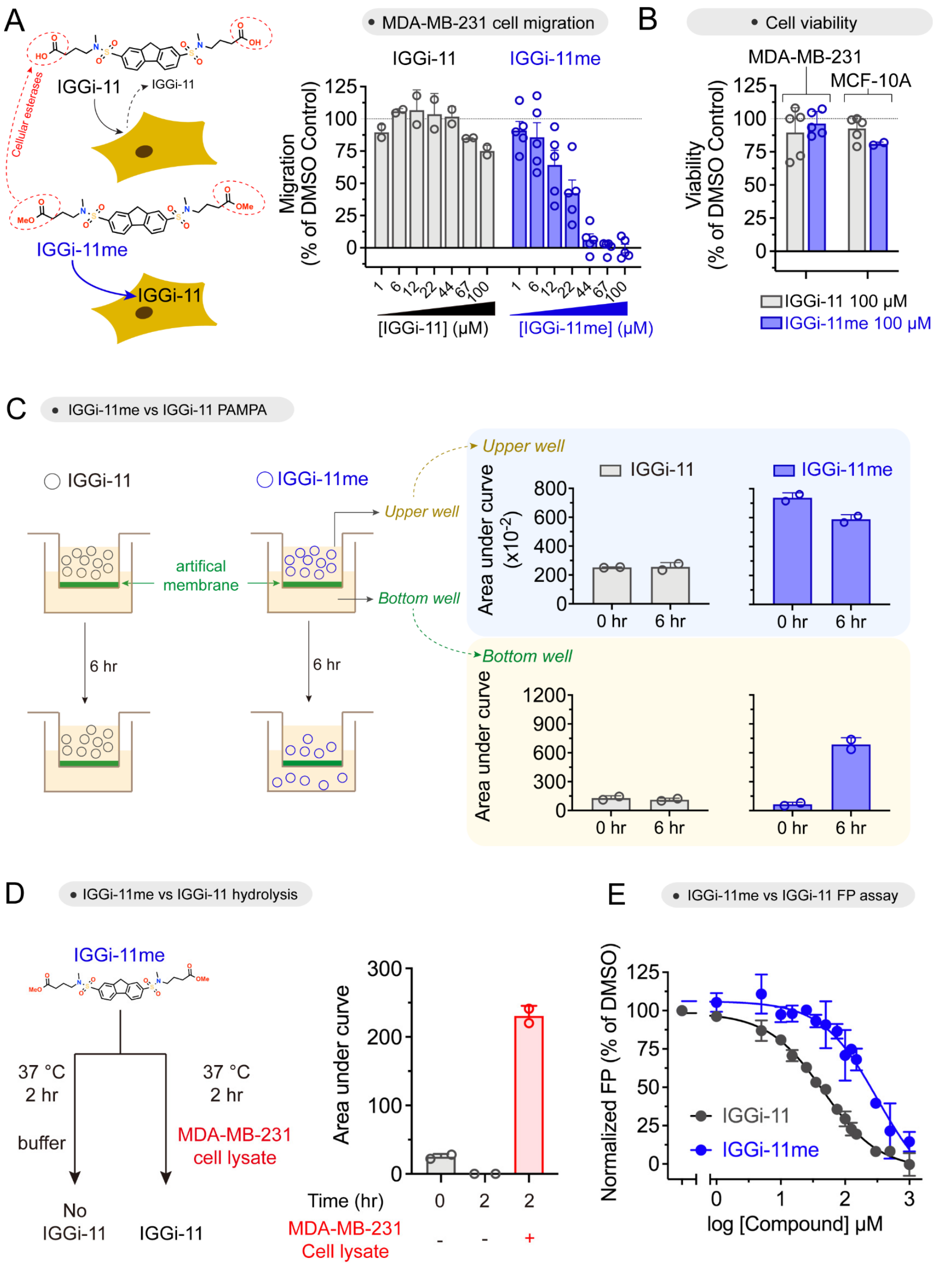


**Figure S7. IGGi-11me is a membrane permeable analog of IGGi-11 that is bioactive in cells. (A)** *Left*, diagram depicting a putative mechanism by which IGGi-11me acts in cells. Esterification of IGGi-11's carboxylate groups to generate IGGi-11me is proposed to increase membrane permeability, whereas action of cellular esterases on IGGi-11me leads to formation of the active compound IGGi-11. *Right*, IGGi-11me, but not IGGi-11, efficiently blocks MDA-MB-231 cell migration. Cell migration was determined using a modified Boyden-chamber assay in the presence of the indicated concentrations of compound (1-100 μM). Results are expressed as % of migration compared to cells treated with DMSO (1 %). Mean ± S.E.M ($N$ = 2-5) **(B)** Neither IGGi-11 nor IGGi-11me affect the viability of MDA-MB-231 or MCF-10A cells. Results are expressed as % of viability of cells treated with 100 μM compound compared to cells treated with DMSO (1 %). Mean ± S.E.M ($N$ = 2-5). **(C)** IGGi-11me displays higher permeability than IGGi-11 in parallel artificial membrane permeability assays (PAMPA). The presence of IGGi-11 or IGGi-11me in the upper and lower wells of the PAMPA assays was determined by LC-MS before and 6 hours after addition of compound to the upper well. Duplicates of one experiment representative of two are presented. **(D)** IGGi-11me is converted to IGGi-11 by cellular esterases. IGGi-11me was incubated in the presence or absence of a cytosolic fraction of MDA-MB-231 cells for 2 hours and the amount of IGGi-11 present in the sample was determined by LC-MS. Duplicates of one experiment representative of two are presented. **(E)** IGGi-11me is less potent than IGGi-11 as an inhibitor of the GIV-Gαi interaction. Binding of GIV to rat Gαi3 was determined by fluorescence polarization in the presence of different concentrations of the indicated compounds. Mean ± S.E.M ($N$ = 3).

FIGURE S8

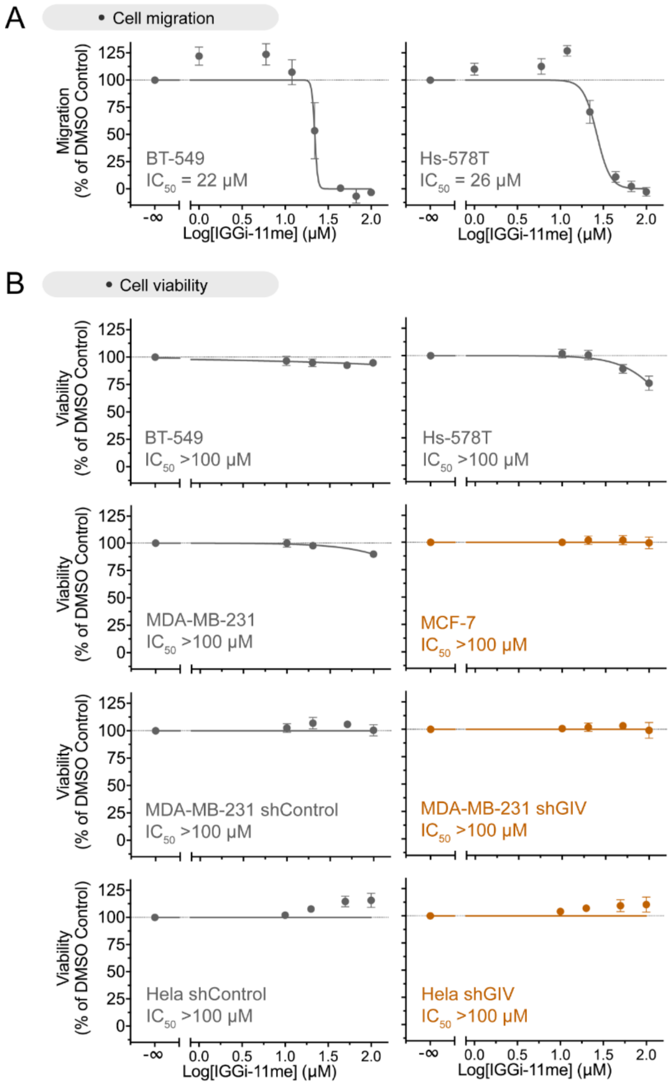


**Figure S8. IGGi-11me blocks cell migration in GIV$^{High}$ cells, but does not affect cell viability in multiple cell lines. (A)** IGGi-11me blocks migration of BT-549 and Hs578T cells, two GIV$^{High}$ cell lines (**Fig. 5**). Cell migration was determined using a modified Boyden-chamber assay in the presence of the indicated concentrations of compound (1-100 µM). Results are expressed as % of migration compared to cells treated with DMSO (1 %). Mean ± S.E.M ($N$ = 3) **(B)** IGGi-11me does not affect the viability of GIV$^{High}$ cells (BT-549, Hs578T, MDA-MB-231, or HeLa cell lines), GIV$^{Low}$ (MCF-7), or GIV-depleted MDA-MB-231 or HeLa cells. Results are expressed as % of viability of cells treated with the indicated concentrations of compound (1-100 µM) compared to cells treated with DMSO (1 %). Mean ± S.E.M ($N$ = 3).

**TABLE S1**

| Structure | IGGi ID | Scaffold | molWeight | Source | IUPAC Name |
|---|---|---|---|---|---|
| 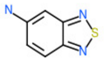 | 1 | Singleton | 151.0 | ChemBridge | 2,1,3-benzothiadiazol-5-amine |
| 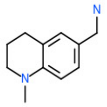 | 2 | G | 176.1 | ChemBridge | (1-methyl-3,4-dihydro-2H-quinolin-6-yl)methanamine |
| 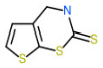 | 3 | Singleton | 187.0 | ChemBridge | 3,4-dihydrothieno[3,2-e][1,3]thiazine-2-thione |
| 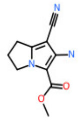 | 4 | Singleton | 205.1 | ChemBridge | methyl 2-amino-1-cyano-6,7-dihydro-5H-pyrrolizine-3-carboxylate |
| 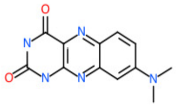 | 5 | E | 257.1 | ChemBridge | 8-(dimethylamino)-1H-benzo[g]pteridine-2,4-dione |
| 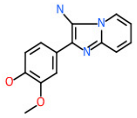 | 6 | Singleton | 255.1 | ChemBridge | 4-(3-aminoimidazo[1,2-a]pyridin-2-yl)-2-methoxyphenol |
| 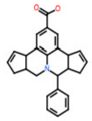 | 7 | G | 369.2 | ChemBridge | 2-phenyl-1-azapentacyclo[10.6.1.03,7.08,19.013,17]nonadeca-5,8,10,12(19),14-pentaene-10-carboxylic acid |
| 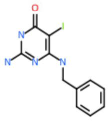 | 8 | F | 342.0 | ChemBridge | 2-amino-4-(benzylamino)-5-iodo-1H-pyrimidin-6-one |
| 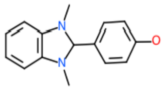 | 9 | Singleton | 240.1 | ChemBridge | 4-(1,3-dimethyl-2H-benzimidazol-2-yl)phenol |
| 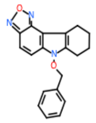 | 10 | Singleton | 319.1 | ChemBridge | 6-phenylmethoxy-7,8,9,10-tetrahydro-[1,2,5]oxadiazolo[3,4-c]carbazole |

| | | | | | |
|---|---|---|---|---|---|
| 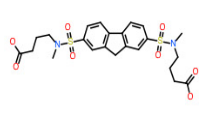 | 11 | C | 524.1 | ChemBridge | 4'-((9H-fluorene-2,7-disulfonyl)bis(methylazanediyl))dibutyric acid |
| 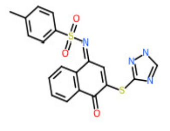 | 12 | B | 410.1 | ChemBridge | 4-methyl-N-[4-oxo-3-(1H-1,2,4-triazol-5-ylsulfanyl)naphthalen-1-ylidene]benzenesulfonamide |
| 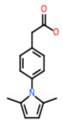 | 13 | I | 229.1 | ChemBridge | 2-[4-(2,5-dimethylpyrrol-1-yl)phenyl]acetic acid |
| 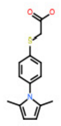 | 14 | I | 261.1 | ChemBridge | 2-[4-(2,5-dimethylpyrrol-1-yl)phenyl]sulfanylacetic acid |
| 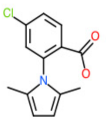 | 15 | I | 249.1 | ChemBridge | 4-chloro-2-(2,5-dimethylpyrrol-1-yl)benzoic acid |
| 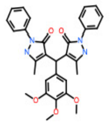 | 16 | A | 526.2 | ChemBridge | 5-methyl-4-[(5-methyl-3-oxo-2-phenyl-1H-pyrazol-4-yl)-(3,4,5-trimethoxyphenyl)methyl]-2-phenyl-1H-pyrazol-3-one |
| 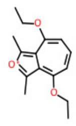 | 17 | Fragment | 247.1 | ChemBridge | (4-ethoxy-1,3-dimethylcyclohepta[c]furan-8-ylidene)-ethyloxidanium |
| 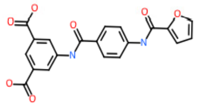 | 18 | J | 394.1 | ChemBridge | 5-[[4-(furan-2-carbonylamino)benzoyl]amino]benzene-1,3-dicarboxylic acid |
| 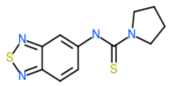 | 19 | D | 264.1 | ChemBridge | N-(2,1,3-benzothiadiazol-5-yl)pyrrolidine-1-carbothioamide |
| 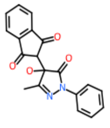 | 20 | A | 334.1 | ChemBridge | 2-(4-hydroxy-3-methyl-5-oxo-1-phenylpyrazol-4-yl)indene-1,3-dione |

| | | | | | |
|---|---|---|---|---|---|
| 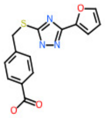 | 21 | Singleton | 301.1 | ChemBridge | 4-[[5-(furan-2-yl)-1H-1,2,4-triazol-3-yl]sulfanylmethyl]benzoic acid |
| 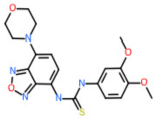 | 22 | D | 415.1 | ChemBridge | 1-(3,4-dimethoxyphenyl)-3-(4-morpholin-4-yl-2,1,3-benzoxadiazol-7-yl)thiourea |
| 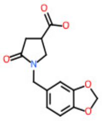 | 23 | Singleton | 263.1 | ChemBridge | 1-(1,3-benzodioxol-5-ylmethyl)-5-oxopyrrolidine-3-carboxylic acid |
| 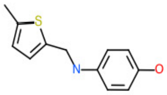 | 24 | J | 219.1 | ChemBridge | 4-[(5-methylthiophen-2-yl)methylamino]phenol |
| 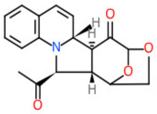 | 25 | Singleton | 311.1 | ChemBridge | (2S,3S,13R,14S)-3-acetyl-17,19-dioxa-4-azapentacyclo[14.2.1.02,14.04,13.05,10]nonadeca-5,7,9,11-tetraen-15-one |
| 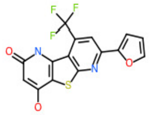 | 26 | C | 352.0 | ChemBridge | 11-(furan-2-yl)-6-hydroxy-13-(trifluoromethyl)-8-thia-3,10-diazatricyclo[7.4.0.02,7]trideca-1(9),2(7),5,10,12-pentaen-4-one |
| 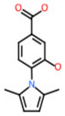 | 27 | I | 231.1 | ChemDiv | 4-(2,5-dimethylpyrrol-1-yl)-3-hydroxybenzoic acid |
| 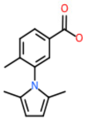 | 28 | I | 229.1 | ChemDiv | 3-(2,5-dimethylpyrrol-1-yl)-4-methylbenzoic acid |
| 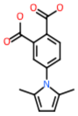 | 29 | I | 259.1 | ChemDiv | 4-(2,5-dimethylpyrrol-1-yl)phthalic acid |
| 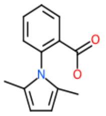 | 30 | I | 215.1 | ChemDiv | 2-(2,5-dimethylpyrrol-1-yl)benzoic acid |

| | | | | | |
|---|---|---|---|---|---|
| 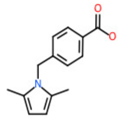 | 31 | I | 229.1 | ChemDiv | 4-[(2,5-dimethylpyrrol-1-yl)methyl]benzoic acid |
| 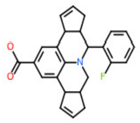 | 32 | G | 387.2 | ChemDiv | 2-(2-fluorophenyl)-1-azapentacyclo[10.6.1.03,7.08,19.013,17]nonadeca-5,8,10,12(19),14-pentaene-10-carboxylic acid |
| 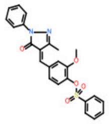 | 33 | A | 448.1 | ChemDiv | [2-methoxy-4-[(3-methyl-5-oxo-1-phenylpyrazol-4-ylidene)methyl]phenyl] benzenesulfonate |
| 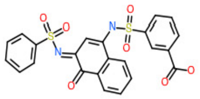 | 34 | B | 496.0 | ChemDiv | 3-[[3-(benzenesulfonamido)-4-oxonaphthalen-1-ylidene]amino]sulfonylbenzoic acid |
| 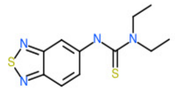 | 35 | D | 266.1 | ChemDiv | 3-(2,1,3-benzothiadiazol-5-yl)-1,1-diethylthiourea |
| 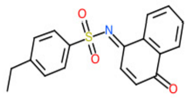 | 36 | B | 325.1 | ChemDiv | 4-ethyl-N-(4-oxonaphthalen-1-ylidene)benzenesulfonamide |
| 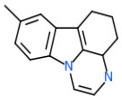 | 37 | Singleton | 224.1 | ChemDiv | 12-methyl-1,4-diazatetracyclo[7.6.1.05,16.010,15]hexadeca-2,9(16),10(15),11,13-pentaene |
| 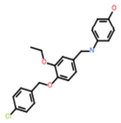 | 38 | J | 383.1 | ChemDiv | 4-[[4-[(4-chlorophenyl)methoxy]-3-ethoxyphenyl]methylamino]phenol |
| 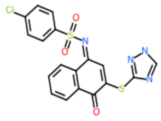 | 39 | B | 430.0 | ChemDiv | 4-chloro-N-[4-oxo-3-(1H-1,2,4-triazol-5-ylsulfanyl)naphthalen-1-ylidene]benzenesulfonamide |
| 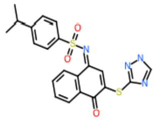 | 40 | B | 438.1 | ChemDiv | N-[4-oxo-3-(1H-1,2,4-triazol-5-ylsulfanyl)naphthalen-1-ylidene]-4-propan-2-ylbenzenesulfonamide |

| | | | | | |
|---|---|---|---|---|---|
| 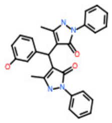 | 41 | A | 452.2 | ChemDiv | 4-[(3-hydroxyphenyl)-(5-methyl-3-oxo-2-phenyl-1H-pyrazol-4-yl)methyl]-5-methyl-2-phenyl-1H-pyrazol-3-one |
| 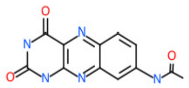 | 42 | E | 271.1 | ChemDiv | N-(2,4-dioxo-1H-benzo[g]pteridin-8-yl)acetamide |
| 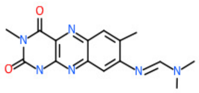 | 43 | E | 312.1 | ChemDiv | N'-(3,7-dimethyl-2,4-dioxo-1H-benzo[g]pteridin-8-yl)-N,N-dimethylmethanimidamide |
| 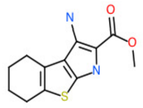 | 44 | C | 250.1 | ChemDiv | methyl 1-amino-5,6,7,8-tetrahydro-3H-[1]benzothiolo[2,3-b]pyrrole-2-carboxylate |
| 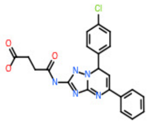 | 45 | Singleton | 423.1 | ChemDiv | 4-[[7-(4-chlorophenyl)-5-phenyl-4,7-dihydro-[1,2,4]triazolo[1,5-a]pyrimidin-2-yl]amino]-4-oxobutanoic acid |
| 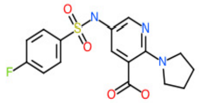 | 46 | B | 365.1 | ChemDiv | 5-[(4-fluorophenyl)sulfonylamino]-2-pyrrolidin-1-ylpyridine-3-carboxylic acid |
| 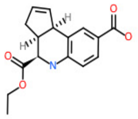 | 47 | H | 287.1 | ChemDiv | (3aS,4R,9bR)-4-ethoxycarbonyl-3a,4,5,9b-tetrahydro-3H-cyclopenta[c]quinoline-8-carboxylic acid |
| 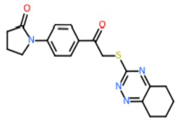 | 48 | Singleton | 368.1 | Asinex | 1-[4-[2-(5,6,7,8-tetrahydro-1,2,4-benzotriazin-3-ylsulfanyl)acetyl]phenyl]pyrrolidin-2-one |
| 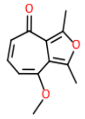 | 49 | Fragment | 204.1 | Asinex | 4-methoxy-1,3-dimethylcyclohepta[c]furan-8-one |
| 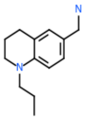 | 50 | G | 204.2 | Asinex | (1-propyl-3,4-dihydro-2H-quinolin-6-yl)methanamine |

| | | | | | |
|---|---|---|---|---|---|
| 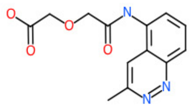 | 51 | Singleton | 275.1 | Maybridge | 2-[2-[(3-methylcinnolin-5-yl)amino]-2-oxoethoxy]acetic acid |
| 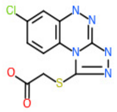 | 52 | Singleton | 297.0 | Maybridge | 2-[(7-chloro-4,5-dihydro-[1,2,4]triazolo[3,4-c][1,2,4]benzotriazin-1-yl)sulfanyl]acetic acid |
| 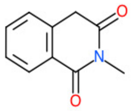 | 53 | Singleton | 175.1 | Maybridge | 2-methyl-4H-isoquinoline-1,3-dione |
| 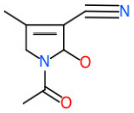 | 54 | Singleton | 166.1 | Maybridge | 1-acetyl-2-hydroxy-4-methyl-2,5-dihydropyrrole-3-carbonitrile |
| 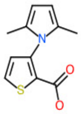 | 55 | I | 221.1 | Maybridge | 3-(2,5-dimethylpyrrol-1-yl)thiophene-2-carboxylic acid |
| 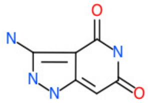 | 56 | Fragment | 166.0 | Maybridge | 3-amino-4-hydroxy-1,5-dihydropyrazolo[4,3-c]pyridin-6-one |
| 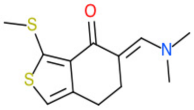 | 57 | Singleton | 253.1 | Maybridge | (5E)-5-(dimethylaminomethylidene)-3-methylsulfanyl-6,7-dihydro-2-benzothiophen-4-one |
| 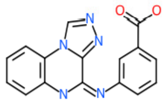 | 58 | Singleton | 305.1 | ChemDiv | 3-([1,2,4]triazolo[4,3-a]quinoxalin-4-ylamino)benzoic acid |
| 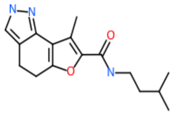 | 59 | C | 287.2 | ChemDiv | 8-methyl-N-(3-methylbutyl)-4,5-dihydro-1H-furo[2,3-g]indazole-7-carboxamide |
| 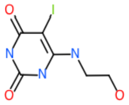 | 60 | F | 297.0 | TimTec | 6-(2-hydroxyethylamino)-5-iodo-1H-pyrimidine-2,4-dione |

| | | | | | |
|---|---|---|---|---|---|
| 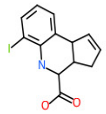 | 61 | H | 341.0 | TimTec | 6-iodo-3a,4,5,9b-tetrahydro-3H-cyclopenta[c]quinoline-4-carboxylic acid |
| 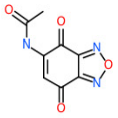 | 62 | Singleton | 207.0 | Ambinter | N-(4,7-dioxo-2,1,3-benzoxadiazol-5-yl)acetamide |
| 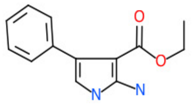 | 63 | Singleton | 230.1 | Ambinter | ethyl 2-amino-4-phenyl-1H-pyrrole-3-carboxylate |
| 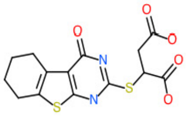 | 64 | C | 354.0 | Ambinter | 2-[(4-oxo-5,6,7,8-tetrahydro-3H-[1]benzothiolo[2,3-d]pyrimidin-2-yl)sulfanyl]butanedioic acid |
| 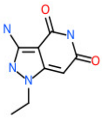 | 65 | Fragment | 194.1 | Ambinter | 3-amino-1-ethyl-4-hydroxy-5H-pyrazolo[4,3-c]pyridin-6-one |
| 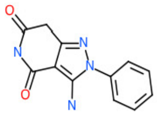 | 66 | A | 242.1 | Ambinter | 3-amino-2-phenyl-7H-pyrazolo[4,3-c]pyridine-4,6-dione |
| 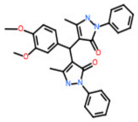 | 67 | A | 496.2 | Ambinter | 4-[(3,4-dimethoxyphenyl)-(5-methyl-3-oxo-2-phenyl-1H-pyrazol-4-yl)methyl]-5-methyl-2-phenyl-1H-pyrazol-3-one |
| 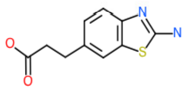 | 68 | Singleton | 222.0 | Sigma | 3-(2-amino-1,3-benzothiazol-6-yl)propanoic acid |
| 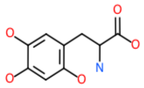 | 69 | Singleton | 213.1 | Sigma | 2-amino-3-(2,4,5-trihydroxyphenyl)propanoic acid |

*of the National Academy of Sciences of the United States of America* 112(9):E937-946.

39. Nemethy G, Gibson KD, Palmer KA, Yoon CN, Paterlini G, Zagari A, Rumsey S, & Scheraga HA (1992) Energy Parameters in Polypeptides .10. Improved Geometrical Parameters and Nonbonded Interactions for Use in the Ecepp/3 Algorithm, with Application to Proline-Containing Peptides. *J Phys Chem-Us* 96(15):6472-6484.
40. Garcia-Marcos M, Kietrsunthorn PS, Wang H, Ghosh P, & Farquhar MG (2011) G Protein binding sites on Calnuc (nucleobindin 1) and NUCB2 (nucleobindin 2) define a new class of G(alpha)i-regulatory motifs. *J Biol Chem* 286(32):28138-28149.
41. Garcia-Marcos M (2021) Complementary biosensors reveal different G-protein signaling modes triggered by GPCRs and non-receptor activators. *eLife* 10.
42. Freissmuth M, Boehm S, Beindl W, Nickel P, Ijzerman AP, Hohenegger M, & Nanoff C (1996) Suramin analogues as subtype-selective G protein inhibitors. *Molecular pharmacology* 49(4):602-611.
43. Parag-Sharma K, Leyme A, DiGiacomo V, Marivin A, Broselid S, & Garcia-Marcos M (2016) Membrane Recruitment of the Non-receptor Protein GIV/Girdin (Galpha-interacting, Vesicle-associated Protein/Girdin) Is Sufficient for Activating Heterotrimeric G Protein Signaling. *J Biol Chem* 291(53):27098-27111.
44. Debnath J, Muthuswamy SK, & Brugge JS (2003) Morphogenesis and oncogenesis of MCF-10A mammary epithelial acini grown in three-dimensional basement membrane cultures. *Methods* 30(3):256-268.
45. Muller RE, Klein KR, Hutsell SQ, Siderovski DP, & Kimple AJ (2010) A homogeneous method to measure nucleotide exchange by alpha-subunits of heterotrimeric G-proteins using fluorescence polarization. *Assay and drug development technologies* 8(5):621-624.
